# Highly potent anti-SARS-CoV-2 multivalent DARPin therapeutic candidates

**DOI:** 10.1101/2020.08.25.256339

**Authors:** Marcel Walser, Sylvia Rothenberger, Daniel L. Hurdiss, Anja Schlegel, Valérie Calabro, Simon Fontaine, Denis Villemagne, Maria Paladino, Tanja Hospodarsch, Alexandra Neculcea, Andreas Cornelius, Patricia Schildknecht, Mirela Matzner, Martin Hänggi, Marco Franchini, Yvonne Kaufmann, Doris Schaible, Iris Schlegel, Chloe Iss, Thamar Looser, Susanne Mangold, Christel Herzog, Dieter Schiegg, Christian Reichen, Filip Radom, Andreas Bosshart, Andreas Lehmann, Micha A. Haeuptle, Alexander Zürcher, Toni Vagt, Gabriel Sigrist, Marcel Straumann, Karl Proba, Niina Veitonmäki, Keith M. Dawson, Christof Zitt, Jennifer Mayor, Sarah Ryter, Heyrhyoung Lyoo, Chunyan Wang, Wentao Li, Ieva Drulyte, Wenjuan Du, H. Kaspar Binz, Leon de Waal, Koert J. Stittelaar, Sarah Taplin, Seth Lewis, Daniel Steiner, Frank J.M. van Kuppeveld, Olivier Engler, Berend-Jan Bosch, Michael T. Stumpp, Patrick Amstutz

**Author notes:** To whom correspondence should be addressed: Michael T. Stumpp. These authors contributed equally to this work (Marcel Walser, Sylvia Rothenberger, Daniel L. Hurdiss, Olivier Engler, Berend-Jan Bosch, Michael T. Stumpp, Patrick Amstutz). Summarizing 125-character sentence: Generation of ensovibep, the first clinical-stage trivalent anti-SARS-CoV-2 therapeutic candidate. DARPin® is a registered trademark owned by Molecular Partners AG.

## Abstract

Globally accessible therapeutics against SARS-CoV-2 are urgently needed. Here, we report the generation of the first anti-SARS-CoV-2 DARPin molecules with therapeutic potential as well as rapid large-scale production capabilities. Highly potent multivalent DARPin molecules with low picomolar virus neutralization efficacies were generated by molecular linkage of three different monovalent DARPin molecules. These multivalent DARPin molecules target various domains of the SARS-CoV-2 spike protein, thereby limiting possible viral escape. Cryo-EM analysis of individual monovalent DARPin molecules provided structural explanations for the mode of action. Analysis of the protective efficacy of one multivalent DARPin molecule in a hamster SARS-CoV-2 infection model demonstrated a significant reduction of pathogenesis. Taken together, the multivalent DARPin molecules reported here, one of which has entered clinical studies, constitute promising therapeutics against the COVID-19 pandemic.

## Introduction

Fighting the COVID-19 pandemic will require coordinated global efforts to maximize the benefits of vaccinations and therapeutics(*1*). Even though vaccine and therapeutic development efforts have progressed considerably, there is, and will be, a remaining medical need for globally accessible therapeutics to treat patients and to protect health care workers, as well as individuals with underlying medical conditions that preclude them from being vaccinated. Neutralizing monoclonal antibodies are expected to be critically important and could be readily available(*2–4*), however they are complex to manufacture and come at a considerable cost. These logistical hurdles may severely limit accessibility, thus preventing an effective global solution(*5*).

DARPin molecules are an emerging class of novel therapeutics that are actively being developed in ophthalmology and oncology, with four molecules at a clinical stage(*6,7*). Here, we report the generation and characterization of the first anti-viral DARPin molecules in the context of the COVID-19 pandemic. DARPin molecules are based on naturally occurring ankyrin repeat motifs. To generate therapeutic DARPin molecules, a pure *in vitro* approach (i.e. selections via ribosome display) is possible and can be carried out in a very short time frame, only requiring the target protein, in this case the SARS-CoV-2 spike protein or subdomains thereof. Hence, therapeutic DARPin molecules can be prepared independently of patient samples or animal immunizations. DARPin molecules can be monovalent and thus monospecific or linked by peptide linkers to form single-chain multivalent or multispecific DARPin molecules with several specificities. Notably, DARPin molecules can be manufactured by microbial fermentation, and thus be potentially available world-wide within a short time due to lower technical requirements to provide large-scale clinical grade material. Additionally, the high heat stability of DARPin molecules offers the prospect of a reduced cold chain for distribution around the globe.

The SARS-CoV-2 spike protein(*8,9*), presented as a metastable prefusion trimer at the viral surface, mediates virus entry into the host cell. The spike protein comprises multiple functional domains: S1, which includes the N-terminal domain (NTD) and the receptor binding domain (RBD) responsible for interaction with the angiotensin-converting enzyme 2 (ACE2) host receptor(*1,8,10,11*), and the S2 domain, which is responsible for virus-host cell membrane fusion via extensive, irreversible conformational changes(*12–14*). This domain composition opens the possibility to target several sites on a single viral protein, leading to multiple mechanisms of inhibition. Such a multi-pronged approach is expected to lead to higher potencies, lower doses, and better protection against potential viral escape mutations.

Here, we present a novel approach using DARPin molecules to simultaneously bind three sites on the trimeric SARS-CoV-2 spike protein. The results reported below describe the development and characterization of monospecific DARPin molecules against distinct domains of the spike protein, the selection process for the most potent monospecific DARPin molecules and, supported by cryo-EM data, their rational combination into highly potent multivalent as well as multispecific DARPin molecules. Furthermore, we demonstrate the protective efficacy of a multivalent DARPin molecule against virus replication and severe disease in a hamster model of COVID-19. We anticipate that antiviral multivalent DARPin molecules have the potential to become an easy-to-deploy antiviral approach for treatment and/or prevention of COVID-19. Based on the results presented here, MP0420 or ensovibep - a multispecific RBD-binding DARPin candidate, is currently being studied in Phase 2 clinical trials.

## Results

### Selection and characterization of monovalent DARPin molecules targeting different regions of the SARS-CoV-2 spike protein

The DARPin technology is based on naïve DARPin libraries(*6*), with a physical diversity of about 10^12^ different monovalent DARPin molecules, allowing the selection of sets of very diverse binding molecules by ribosome display(*15,16*), the method of choice when dealing with libraries of such large diversities. DARPin libraries are based on a consensus design approach using hundreds of ankyrin repeat protein sequences of the human and mouse genome(*17*). An overview of the entire generation process of anti-SARS-CoV-2 spike protein binding DARPin molecules is shown in Figure 1. To obtain individual DARPin molecules binding to distinct domains of the SARS-CoV-2 spike protein and potentially inhibiting viral cell entry, we focused on generating DARPin molecules binding to the receptor binding domain (RBD), the S1 N-terminal domain (NTD) or the S2 domain(*18*). After four ribosome display selection rounds (Figure 1A), we further enriched for the most potently binding DARPin molecules through screening of 3’420 *E. coli* cell extracts overexpressing individual DARPin molecules by homogeneous time-resolved fluorescence (HTRF) assays for binding to different spike protein domains (Figure 1B). Based on binding and ACE2 inhibition profiles obtained in HTRF, which allowed mapping of monovalent DARPin molecules to different spike domains, 380 DARPin molecules were selected to be expressed in 96-well format and purified to homogeneity. DARPin molecules were further characterized for antiviral potency in a VSV-pseudovirion neutralization assay (PsV NA) as well as biophysically by size exclusion chromatography (SEC), Sypro-Orange thermal stability assessment(*19*), ProteOn surface plasmon resonance (SPR) target affinity assessment, and ELISA, to orthogonally evaluate target binding (Figure 1C and D). In parallel to the characterization of the 380 monovalent DARPin molecules, 6 monovalent DARPin molecules of known spike domain specificity were used to randomly assemble a set of 192 tri-specific DARPin molecules (Figure 1E). The antiviral potencies, determined in a PsV-NA screening assay, of these randomly combined tri-specific DARPin molecules provided valuable information on the most potent tri-specific combinations and formats. Based on the combined data for the 380 monovalent DARPin molecules, 11 of them with low pM to low nM affinities, excellent biophysical properties, diversities in amino acid sequences as well as binding for various SARS-CoV-2 spike protein domains (Supplementary Table 1 and Supplementary Figure 1) were selected for the rational generation of 22 multivalent DARPin molecules described below (Figure 1F). After detailed characterization of these 22 multivalent DARPin candidates, systemic exposure was assessed in mice and in Syrian golden hamsters for the most promising multivalent DARPin candidates (Figure 1G). The multivalent DARPin candidate with the longer systemic half-life was evaluated for SARS-CoV-2 protection in a Covid-19 Syrian golden hamster model (Figure 1H).

**Figure 1:**
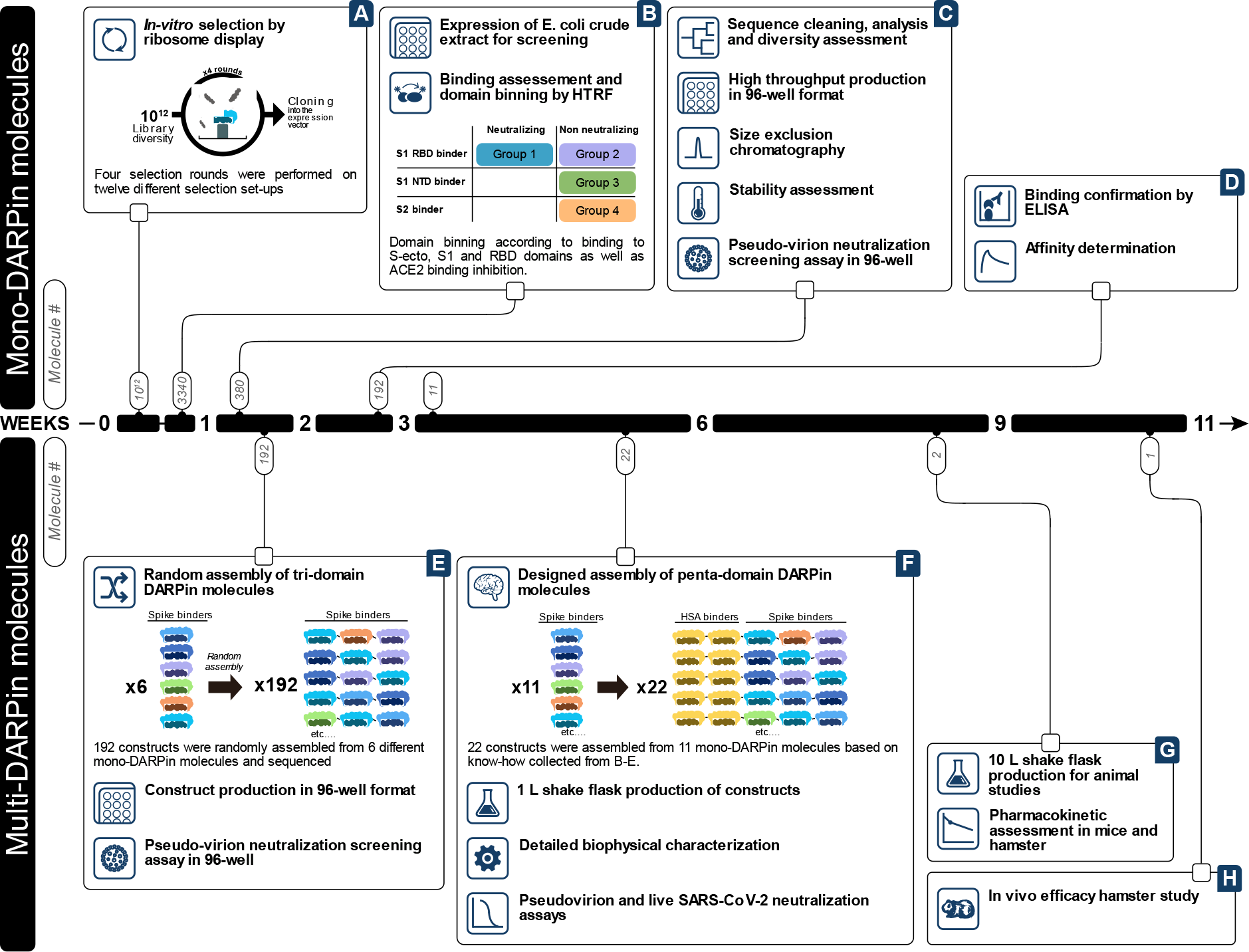
Process overview for the generation of anti-SARS-CoV-2 multivalent DARPin molecules. Upper panel, generation and evaluation of monovalent DARPin molecules. Lower panel, assembly and deep-characterization of multivalent DARPin molecules.

### Rational design of multivalent DARPin molecules targeting the SARS-CoV-2 spike protein

We aimed to increase virus neutralizing potency through molecular linkage of monovalent DARPin molecules leading to avidity effects. Using the 11 selected mono-specific DARPin molecules, a total of 22 multivalent DARPin molecules were generated and characterized in detail, each comprising 3 monovalent DARPin molecules against various epitopes of the spike protein as well as two monovalent DARPin molecules binding to human serum albumin (HSA), which have been previously shown to confer long half-life to other DARPin molecules in animals and humans(*7*) (Supplementary Table 2). Each of the 22 multivalent DARPin molecules contained at least one RBD-binding domain since preventing ACE2 receptor interaction is assumed to be the strongest point of interference with virus entering the host cell. Multivalent DARPin molecules were designed to contain several binding modules to RBD (multi-RBD DARPin molecules) or to several distinct domains (multi-mode DARPin molecules). Based on profiling of 22 multivalent DARPin molecules for their biophysical properties and antiviral potency (Supplementary Table 2), we selected two multivalent DARPin molecules, representative for the two different design strategies (i.e. Multi-RBD-DARPin-Candidate, MR-DC, and Multi-Mode-DARPin-Candidate, MM-DC) for further analysis (Figure 2b). DNA sequencing of the individual monovalent DARPin components of MR-DC revealed a similar, but not identical, arrangement of amino acids, suggesting that they share a common mechanism of binding to the RBD. In contrast, the individual components of MM-DC exhibited a high level of sequence diversity, consistent with their different targeting mechanisms (Figure 2c). Neutralization potency for MR-DC and MM-DC was 56 and 36-fold higher in PsV NA, respectively, relative to the most potent neutralizing individual monovalent DARPin that was used for the design of the two types of multivalent DARPin molecules (Figure 2d-e). In addition, IC_50_ values for potency determination in a VSV-SARS-CoV2 pseudotype neutralization assay for MR-DC and MM-DC was similar or superior to that of three earlier described potent neutralizing antibodies(*20,21*). Neutralization assays with the infectious SARS-CoV-2 yielded IC_50_ values of 12 pM for MR-DC (1 ng/ml) and 80 pM for MM-DC (7 ng/ml), respectively (Figure 2f). Still, despite slightly higher potencies displayed by some other screened multi-valent DARPin candidates (e.g. M6, M7, M20 and M22), candidates MR-DC and MM-DC were selected as lead candidates to provide the highest diversity within their paratopes, and potentially the best possible protection against viral escape mutations. In addition, both multivalent DARPin molecules could effectively neutralize pseudotype viruses carrying spike proteins containing natural occurring polymorphisms, including the frequently occurring D614G mutation (Supplementary Table 3). The extremely high neutralization potencies are key for SARS-CoV-2 treatment in particular in a prophylactic setting where very low virus titers at the beginning of the infection are expected.

**Figure 2:**
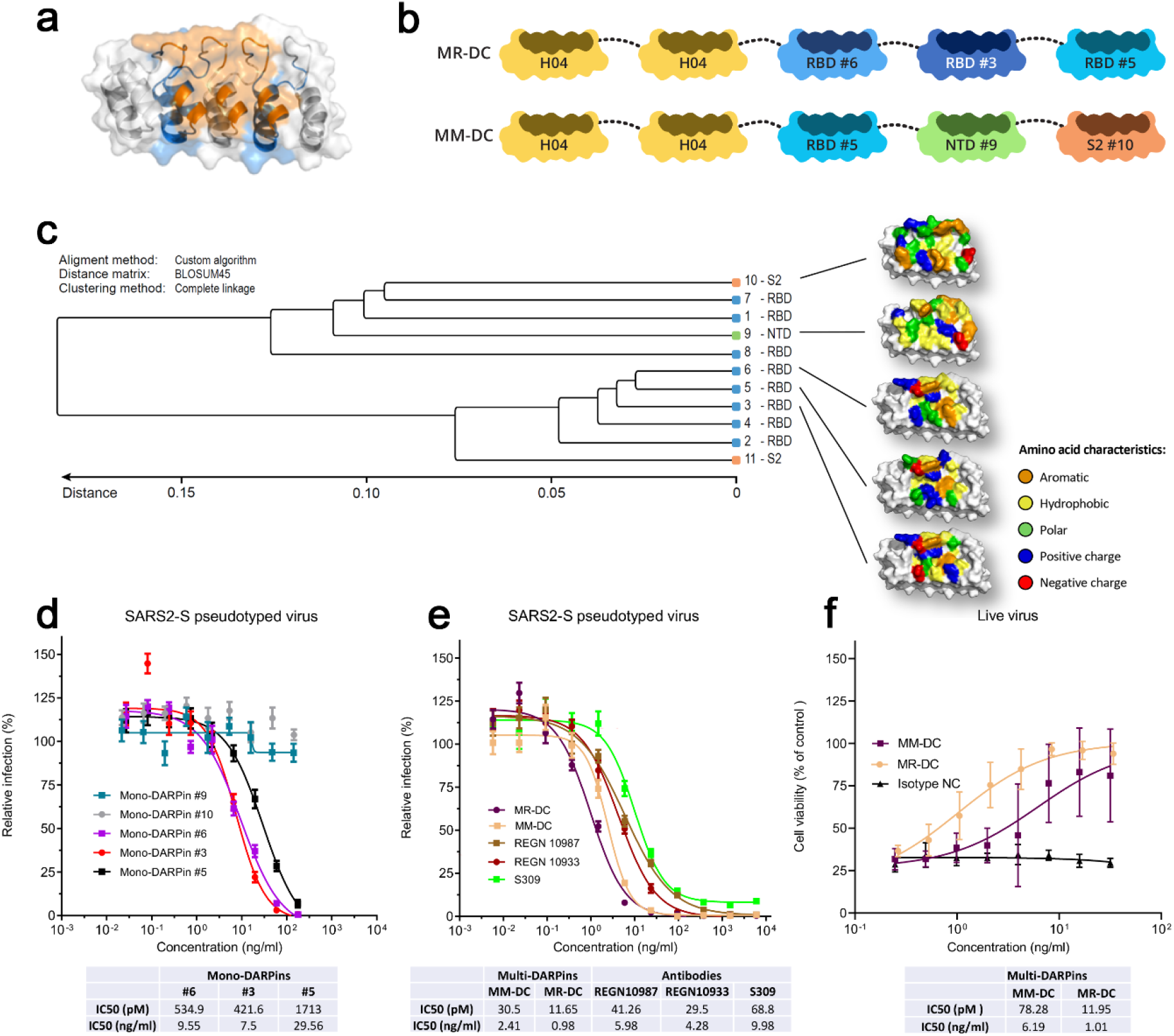
Architecture, diversity and *in vitro* potency of two multivalent DARPin molecules. a) Ribbon structure and semi-transparent surface representation of a monovalent DARPin molecule. The designed ankyrin repeats are colored alternatingly in white and blue for the five repeats. The randomized residues present in the rigid target binding surface are colored orange. b) Schematic overview of the MR-DC and MM-DC constructs. Protein linkers are depicted as gray dashed lines and the half-life extending human serum albumin binding monovalent DARPin (H) is colored yellow. c) Sequence family tree illustrating the sequence diversity amongst the 11 monovalent DARPin molecules chosen for the generation of multivalent DARPin molecules. Surface representations of five Monovalent DARPin molecules binding to the RBD, NTD or S2 are shown, with the amino acid residues in the binding surface colored according to their biophysical characteristics as indicated. d, e) DARPin-respectively antibody-mediated neutralization of infection of luciferase-encoding VSV particles pseudotyped with the SARS-CoV-2 spike protein. Pseudotype VSV particles pre-incubated with monovalent DARPin molecules (d), multivalent DARPin molecules (MM-DC and MR-DC) or control antibodies (e) at the indicated concentrations were used to infect VeroE6 cells. Luciferase activities in cell lysates were determined at 24 h post transduction to calculate infection (%) relative to mock-treated virus controls. The average ± SD from two independent experiments performed in sextuplicate is shown. f) Multivalent DARPin-mediated cell protection of SARS-CoV-2 infection. SARS-CoV-2 pre-incubated with multivalent DARPin molecules MM-DC or MR-DC, or with isotype negative control (isotype NC) at the indicated concentrations were used to infect VeroE6 cells. Cell viability was determined using the CellTiter-Glo luminescent cell viability assay and represented as relative (%) to mock infected cells. Half-maximal inhibitory concentration values (IC_50_) of DARPin molecules and antibodies are indicated in the lower panel tables (d-f).

### Cryo-EM analysis of monovalent DARPin molecules

To gain further insights into the mode of inhibition and binding, three individual monovalent DARPin molecules targeting RBD, S1-NTD, or S2, were subjected to cryo-electron microscopy (cryo-EM) analysis in complex with the trimeric spike ectodomain. In each case, few intact spikes remained following incubation with the monovalent DARPin molecules, particularly with the S2 binder (Supplementary Figure 2). When spike ectodomains were incubated with RBD-binding monovalent DARPin molecule #3 for 15 seconds prior to vitrification, 3D classification revealed that 65% of the intact S ectodomains were in the closed conformation, 20% had two RBDs in the open conformation and 15% had all three RBDs in the open conformation (Supplementary Figure 3a). For the open RBD classes, additional density, consistent with the size of a monovalent DARPin molecule, was present on the RBD receptor binding ridge (RBR). When the incubation time was increased to 60 seconds, 66% of S ectodomains had three monovalent DARPin-bound RBDs in the open conformation (Supplementary Figure 3b). Interestingly, 18% of the S ectodomains had two DARPin-bound RBDs in the open conformation and one trapped in a partially-closed conformation (Supplementary Figures 3b and 4). These results demonstrate that monovalent DARPin #3 binding prevents closure of the RBD through a previously described ratcheting mechanism(*18*). 3D refinement of the fully open class, from the 60 second incubated sample, produced a 4.2 Å global resolution map (Figure 3a and Supplementary Figure 3c-e). The resolution of the map was sufficient to unambiguously assign the pose of RBD-binding monovalent DARPin molecule #3, which binds perpendicular to the RBD receptor binding motif (RBM), with its N-terminus orientated toward the spike three-fold symmetry axis (Figure 3a). The concave DARPin binding surface covers the RBR and overlaps with the ACE2 binding site (Figure 3b). Based on the cryo-EM data, molecular docking experiments were performed. The top scoring model indicated that the interface area is ∼700 Å^2^ and that key epitope residues are F456, Y473, F486, N487 and Y489, which putatively form an interface of hydrophobic interactions and hydrogen-bonds with the DARPin molecule #3 (Supplementary Figure 5). Taken together, these data show that RBD-binding monovalent DARPin molecule #3 inhibits SARS-CoV-2 through receptor functional mimicry, facilitating fusogenic conformational rearrangements of the spike protein. This mechanism of inhibition was also observed for a SARS-CoV neutralizing antibody, S230(*18*), and more recently for a SARS-CoV-2 neutralizing antibody, C105(*22*). Based on our cryo-EM structure, molecular modelling was used to demonstrate that a linker, used in the multivalent DARPin format, would permit simultaneous binding of three RBD-targeting monovalent DARPin molecules (Figure 3c).

**Figure 3:**
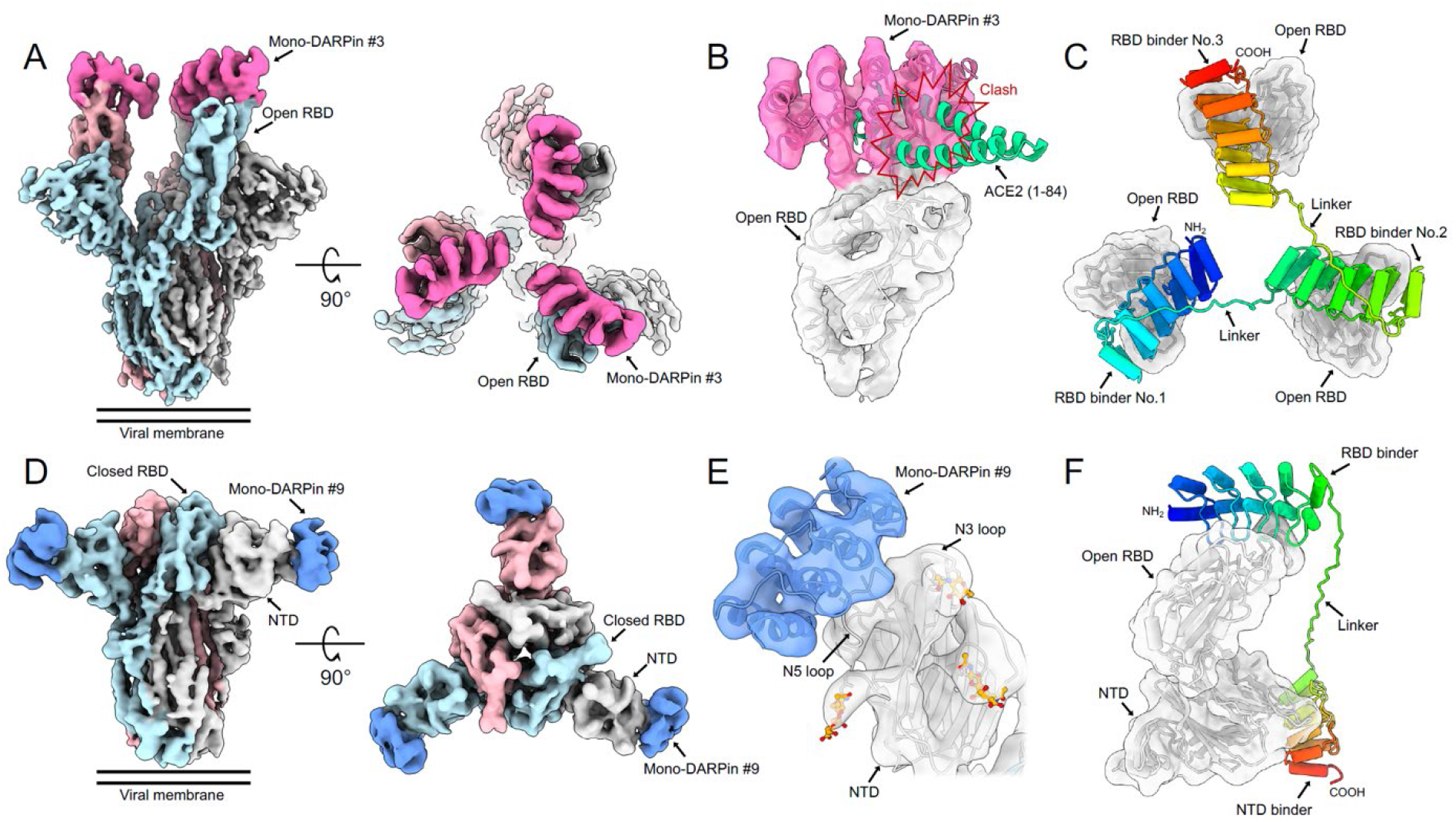
Cryo-EM analysis of monovalent DARPin molecules. A) Cryo-EM density for the SARS-CoV-2 spike ectodomain in complex with the RBD targeting monovalent DARPin #3, shown as two orthogonal views. The DARPin density is colored magenta and the three spike protomers are colored light blue, grey and pink. B) Zoomed in view of a single DARPin #3-bound RBD with the cryo-EM density shown semi-transparent. The atomic coordinates for the fitted fully open spike (PDB ID: 6XCN) and the DARPin homology model are overlaid. The atomic coordinates for residues 1-84 of the RBD bound ACE2 (PDB ID: 6M0J), colored green, is superimposed. C) Proposed model of three covalently linked RBD-targeting monovalent DARPin molecules shown in a rainbow color scheme from the N terminus (blue) to the C terminus (red). D) Cryo-EM density for the SARS-CoV-2 spike ectodomain in complex with the NTD targeting monovalent DARPin #9, shown as two orthogonal views. The mono-DARPin #9 density is colored blue and the three spike protomers are colored light blue, grey and pink. E) Zoomed in view of the DARPin molecule #9 bound to the NTD with the cryo-EM density shown semi-transparent. The atomic coordinates for the fitted fully closed spike (PDB ID: 6ZGE) and the DARPin homology model are overlaid and colored grey and blue, respectively. The N3 and N5 loops are labelled and glycans are shown in stick representation and colored orange. F) Proposed model of the covalently linked NTD and RBD targeting monovalent DARPin molecules, shown in a rainbow color scheme from the N terminus (blue) to the C terminus (red).

Following a 15 second incubation with the S1-NTD-binding monovalent DARPin molecule #9, 2D classification already revealed clear additional density bound to the spike NTD (Supplementary Figure 2). Subsequent 3D classification revealed that most of the DARPin-bound spikes were in the closed conformation, and 19% had one RBD in open conformation (Supplementary Figure 6a). 3D refinement of the fully closed class produced an 8 Å global resolution map (Figure 3d and Supplementary Figure 6b-c), sufficient to dock and assign the pose of the bound DARPin molecule (Figure 3d). This monovalent DARPin molecule binds to the most distal region of the NTD, which is not resolved in the majority of available spike structures(*13,23*). However, several spike structures with nearly completely modelled NTDs were recently reported(*24,25*). The better resolved NTD loops in these structures allowed us to further narrow down the DARPin epitope, indicating that the concave binding surface is ideally situated to interact with the N5 loop, encompassing residues 246-258(*26*) (Figure 3e). A recently described monoclonal antibody, 4A8(*26*), also targets the NTD, involving an extensive hydrophilic interaction network with the N3 loop and, to a lesser extent, the N5 loop. In contrast, molecular docking experiments suggest monovalent DARPin molecule #9 interacts with the NTD primarily with N5 loop residues 248-252 (Supplementary Figure 7), with an interface area of ∼600 Å^2^ and involving both hydrophilic and hydrophobic interactions. Based on our cryo-EM structures, molecular modelling was used to demonstrate that a linker, used in the multivalent DARPin format, would permit simultaneous binding of the NTD and RBD-targeting monovalent DARPin molecules (Figure 3f).

Cumulatively, our structural analysis allowed us to generate models of the MR-DC and MM-DC molecules in contact with the spike protein using 3D molecular modeling tools (Supplementary Figure 8). In both cases, the half-life extension modules have sufficient space to bind HSA. Within the distance limited by the linker length, we identified a potential binding site for the S2 binder, suggesting that simultaneous binding to the spike protein is feasible. However, structural analysis of S ectodomains incubated with the S2 binder #10 did not reveal any unambiguous density for the bound monovalent DARPin. The low number of intact ectodomains remaining after incubation with monovalent DARPin #10 suggests that, compared to monovalent DARPin molecules #3 and #9, the S2 binder has the greatest destabilizing effect on the structural integrity of the S ectodomains (Supplementary Figure 2).

### In vivo antiviral efficacy of a multivalent DARPin in a SARS-CoV-2 hamster infection model

In order to assess the *in vivo* antiviral efficacy of our multivalent DARPin molecule MR-DC, a hamster Covid-19 disease model was performed to study the potential for preventing SARS-CoV-2 related disease (Figure 4a). Syrian golden hamsters (6 females per group) were treated with a single intraperitoneal dose of one multivalent DARPin molecule, MR-DC, using either 16 µg, 160 µg, 1600 µg (per animal; average body weight of ∼160g), or with placebo, 24 h prior to intranasal infection with SARS-CoV-2. Readouts included observation of macroscopic assessment of tissues, histopathology, body weight and virus titers. Dose-dependent reduction in tissue damage, immune response to infection, body weight loss, virus titers in throat, nasal turbinates and lung tissue was observed, indicating significant anti-viral activity for the 1600 µg dose and a trend for protection for the 160 µg dose under the conditions of the hamster model (Figure 4b-g).

**Figure 4:**
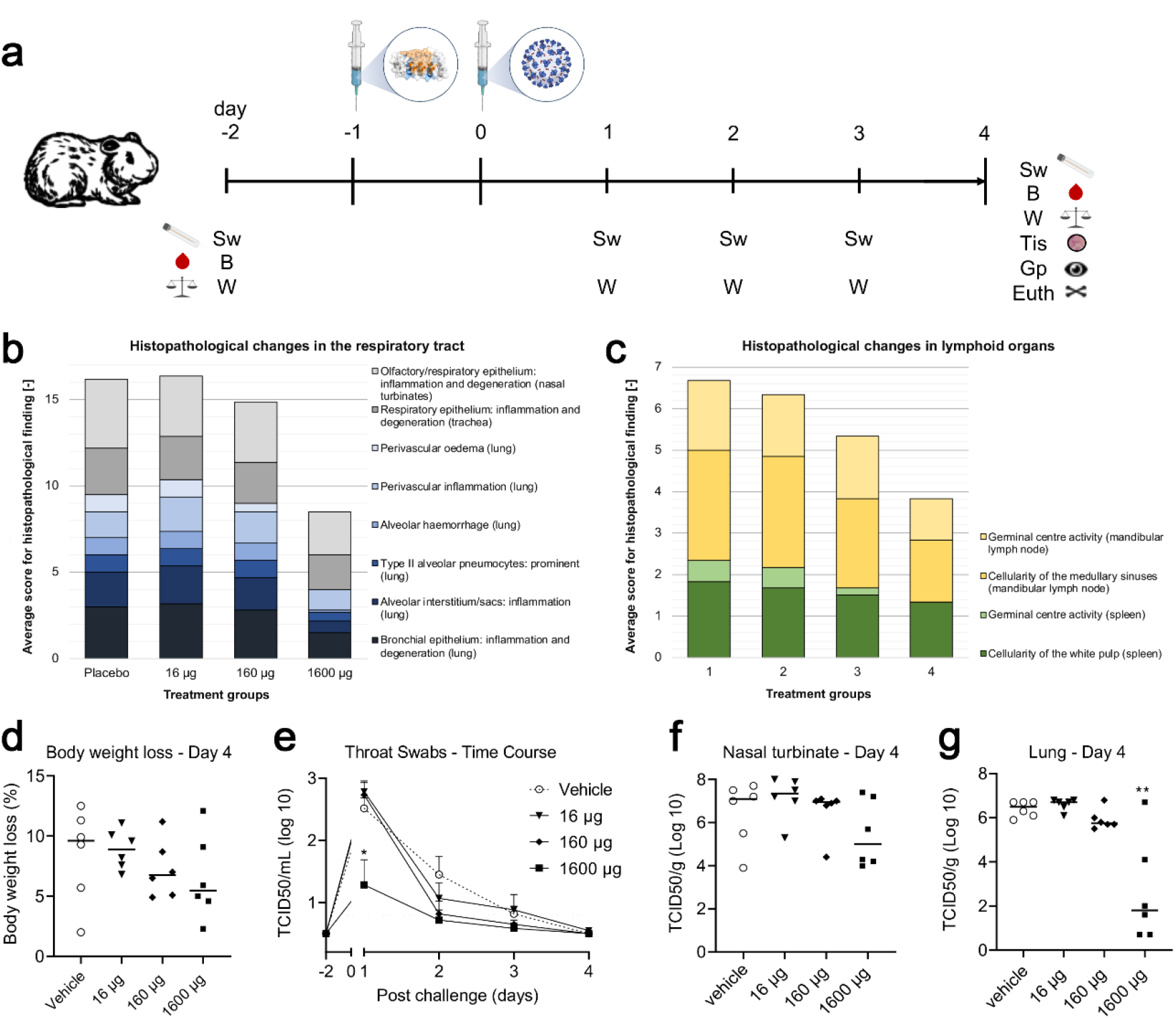
In-vivo efficacy of MR-DC in treating SARS-CoV-2 infection in a preventive Syrian golden hamster model. a) study design: six animals were used per dose group. Generally high variations were observed in control and treatment groups. At Day -2, body weight (W) was measured, blood (B) was taken, and the first throat swab (Sw) performed. Animals were euthanized on Day 4 and tissue (Tis) were taken and gross pathology (Gp) was performed. (b) Histopathological changes in the respiratory tract (lung - blue bars, trachea and nasal turbinates – grey bars). (c) Histopathological changes in lymphoid organs (spleen - green bars, mandibular lymph nodes - yellow bars). A trend to dose-dependent reduction of body-weight loss at day 4 (d), virus titers post challenge for throat swabs (e), nasal turbinate (f) and significant reduction of viral loads in the lung at 1600 µg (10 mg/kg), with two animals below detection limit, as well as a trend to dose-dependent reduction of viral loads at 160 µg (1 mg/kg) (g). Statistically significant differences between Placebo and 1600 µg group: *T-test; P value = 0.0183 / **T-test; P value = 0.0031.

Histopathology results: Intranasal infection with SARS-CoV-2 induced epithelial inflammation and degeneration of the respiratory tract from the nasal turbinates, through the trachea to the bronchi/bronchioles of the lung. More specifically, the inflammation comprised a mixed inflammatory cell infiltrate of lymphocytes, macrophages, plasma, cells and granulocytes which was variably accompanied by epithelial degeneration, regeneration, disorganization, single cell necrosis and inflammatory exudates in the bronchial lumen and nasal cavity. In the lung of more severely affected animals the inflammation extended from the bronchial epithelium to adjacent alveolar interstitium, alveolar sacs and blood vessels (with or without perivascular edema); prominent Type II alveolar pneumocytes and variable hemorrhage were also variably present within the lesion. These findings were most severe in SARS-CoV-2 infected animals receiving the vehicle and the lowest dose of 16ug multivalent DARPin MR-DC. While there was some reduction in the severity of the lesion in infected animals given 160 ug, there was a significant improvement in animals treated with the highest dose of 1600 ug (Figure 4b; Supplementary Figure 9).

SARS-CoV-2 infection was accompanied by a reactive change in lymphoid organs consistent with immune activation. Activation of germinal centers was accompanied by increased cellularity of the white pulp in the spleen and increased cellularity of the medullary sinuses (macrophages and plasma cells) in the mandibular lymph node. Again, these findings were generally more apparent in infected animals receiving the vehicle, with a dose-related decrease in severity in MR-DC-treated groups, particularly in animals treated with the highest dose of 1600 ug (Figure 4c; Supplementary Figure 9). The pharmacokinetics for MR-DC, injected at 160 µg and 1600 µg per animal, was evaluated in an independent study in non-infected Syrian golden hamsters, proving systemic exposure for the duration of the in-life phase of this hamster PD model (Supplementary Figure 10).

## Discussion

Multiple strategies to combat the COVID-19 pandemic are urgently needed. Next to preventive vaccination approaches, monoclonal antibodies are showing therapeutic promise, based on highly potent virus inhibition and encouraging animal and clinical efficacy(*27–29*). However, due to the global need for COVID-19, manufacturing is expected to become a major bottleneck for vaccines and antibodies. A number of alternative molecules are being developed to complement and partially overcome this limitation of antibodies. Here we describe the generation of DARPin molecules - one prominent alternative to antibodies(*30*) amongst others(*31–34*) - that bind the SARS-CoV-2 spike protein. We tested a library of one trillion DARPin molecules and identified multiple DARPin molecules with different functionalities and binding specificities. By molecular linkage of individual DARPin molecules, we developed multivalent DARPin molecules with low picomolar neutralizing activity and demonstrated their protective efficacy against SARS-CoV-2 infection in a hamster model. In particular, reduced lung tissue damage and reduced virus replication in the lower and upper respiratory tract were observed, the latter also being important for reducing virus shedding and transmission. The most advanced of those multivalent DARPin molecules, MP0420 or ensovibep, is currently being studied in Phase 2.

The most potent neutralizing monovalent DARPin molecules were found to target the RBD, blocking the spike-ACE2 interaction necessary for infection. This finding is congruent with the identified epitopes of potent neutralizing antibodies obtained from COVID-19 patients(*20,35–39*). Thus, the *in vitro* approach using DARPin molecules independently confirms that the ACE2 interaction site on the SARS-CoV-2 spike protein is one of the most vulnerable sites to block virus infection in cell culture.

The single-chain binding domain nature of DARPin molecules facilitated the design of multiparatopic and multispecific DARPin molecules with greatly increased neutralization potencies. Virus neutralization capacity increased substantially when three DARPin molecules were linked - relative to the individual modules - likely due to both, increased avidity when binding to the trimeric spike proteins as well as multiple spike domains. A similar strategy with comparable outcome has been recently described for nanobodies. The observed *in vitro* neutralization capacity of the multivalent DARPin molecules in the low picomolar range is similar to or even outcompetes the neutralizing capacities of monoclonal antibodies. The protection against SARS-CoV-2 infections demonstrated in a hamster model at 1600 mg per animal (10 mg/kg) and the partial protection at 160 mg (1 mg/kg) is in the range reported for monoclonal antibodies(*27,31,37,40,41*).

The multivalent DARPin molecules are expected to retain potency even if the spike protein of SARS-CoV-2 should mutate considerably in the future, especially when multiple modes of action are addressed. Evaluation of the impact of a panel of reported mutations in the SARS-CoV-2 spike protein revealed no loss of neutralizing capacity of our lead candidates in a PsV NA and a set of emerging variants and spike protein mutations were recently reported elsewhere(*42*). Although it remains to be determined if additional mutations will impact the neutralization potencies of the multivalent DARPin molecules, we expect multivalent DARPin molecules to retain high potency in case the SARS-CoV-2 spike protein should mutate in the future.

DARPin molecules(*30*) have shown good promise for both ophthalmology and oncology patients. Within oncology, MP0250 is the DARPin molecule(*7*) with most systemic data available to date: MP0250 is a dual inhibitor of VEGF and HGF and is being developed for multiple myeloma in combination with small molecules. MP0250 shows a long systemic half-life of about 2 weeks in human, low immunogenicity potential, and has been given to individual patients for more than two years. All 42 patients in a phase I clinical trial maintained systemic exposure after repeated dosing while 2 patients, out of these 42 patients, showed elevated anti-drug antibodies which did not affect MP0250 exposure. Here, we demonstrate prophylactic protection from SARS-CoV-2 infection by a multivalent DARPin molecule in a hamster model. Reported systemic exposures, achieved with human serum albumin binding DARPin molecules, appear comparable to the half-lives reported for monoclonal antibodies. Due to the lack of an immune stimulating Fc-fragment, we envision no generation of antibody dependent enhancement (ADE) effects, a potential side-effect of monoclonal antibodies in patients with inflamed lungs(*43–46*). This hypothetical differentiation, when compared to monoclonal antibody treatments including immune stimulating Fc-fragments, is currently under clinical investigation and remains to be verified.

Generally, DARPin molecules demonstrate excellent temperature stability, which might enable alternative routes of administration, such as inhalation described for other highly stable protein scaffolds(*47,48*). The simple molecular architecture of multivalent DARPin molecules allows manufacturing in bacterial expression systems at very high yields which leads to cost-effective and scalable production of antiviral biologicals at industrial quantities (supplementary Table 5). Consequently, global access to this novel class of therapeutics may be highly additive to monoclonal antibody approaches and thus contribute to overcoming the COVID-19 pandemic(*49*). We anticipate that the presented workflow for DARPin development can be applied for any future emerging (corona)virus. We have proven that high-affinity binding and potently neutralizing DARPin molecules can be developed in a matter of weeks, without the requirement of immunization of animals or access to patient materials. Such fast track development strategies of picomolar inhibitors are critical to raise preparedness level towards novel pandemic viruses.

## Data availability

The EM density maps for the SARS-CoV-2 spike ectodomain in complex with monovalent DARPin #3 (state 1 and state 2), and monovalent DARPin #9 have been deposited to the Electron Microscopy Data Bank under the accession codes EMD-11953, EMD-11954 and EMD-11956, respectively. The monovalent DARPin and multivalent DARPin sequences, and pseudo-atomic models derived from molecular docking experiments, are available (by contacting the corresponding author) for research purposes only under an MTA, which allows the use of the data for non-commercial purposes but not their disclosure to third parties.

## Acknowledgements

We would particularly like to thank those colleagues at Molecular Partners who are not included in the author list and are currently advancing the program through the clinic. Additionally, we would like to thank William Lee, former board member of Molecular Partners - and the Virology group at Gilead Sciences for their helpful input. The authors also thank Dr. Gert Zimmer for the gift of the recombinant VSV (Institute of Virology and Immunology (IVI), CH-3147 Mittelhäusern, Switzerland, Department of Infectious Diseases and Pathobiology, Vetsuisse Faculty, University of Bern, CH-3012 Bern, Switzerland). The expression plasmid for the SARS-CoV-2 spike protein was kindly provided by Dr. Giulia Torriani and Dr. Isabella Eckerle (Department of Medicine, University of Geneva, Switzerland). SR & JM were supported by Swiss Federal Office for Civil Protection (Grants Nr. 353008564/Stm, 353008218/Stm, and 353008560/Stm to Olivier Engler and Stefan Kunz).

We would like to thank for the supply of the two 2019-nCoV strains: i) 2019-nCoV/IDF0372/2020 provided by the National Reference Centre for Respiratory Viruses hosted by Institut Pasteur (Paris, France) and headed by Dr. Sylvie van der Werf as well as the human sample from which strain 2019-nCoV/IDF0372/2020 was isolated, provided by Dr. X. Lescure and Pr. Y. Yazdanpanah from the Bichat Hospital.

D.L.H. is funded from by the European Union’s Horizon 2020 research and innovation program under the Marie Skłodowska-Curie grant agreement (No 842333) and holds an EMBO non-stipendiary long-term Fellowship (ALTF 1172-2018). Cryo-EM data processing was carried out on the Dutch national e-infrastructure with the support of SURF Cooperative.

## Funding

All studies were funded by Molecular Partners AG, Switzerland.

## Author contributions

M.W., S.R., D.L.H., V.C., C.R., S.F., A.Z., N.V., K.M.D., C.Z., M.A.H., L.d.W., K.J.S., F.J.M.v.K., O.E., B.-J. B., M.T.S.; P.A.: conceptualized and designed experiments; M.W., S.R., D.L.H., A.S., D.V., M.P., T.H., A.N., A.C., P.S., M.M., M.H., M.F., Y.K., I.S., C.I., T.L., S.M., C.H., D.S., A.B., A.L., T.V., G.S., K.P., M.S., J.M., S.Ry., S.T., H.L., C.W., W.L., I.D.: performed laboratory experiments; D.L.H., C.R, F.R.: performed computational work; M.W., S.R., D.L.H., H.K.B., S.L., S.F., F.J.M.v.K., O.E., B.-J. B., M.T.S., P.A. drafted the manuscript. S.L., D.S., M.T.S., P.A. acquired funding. All authors reviewed the manuscript.

## Competing interests

Molecular Partners authors own performance share units and/or stock of the company. HKB owns stock of the company.

## Materials and Methods

### SARS-CoV-2 spike proteins variants used

Proteins used for selections comprised SARS-CoV-2 S protein ectodomain (SARS2-Secto-d72-GCN4-Streptag; University of Utrecht), SARS-CoV-2 S protein (S1+S2 ECT, His-tag; Sinobiological 40589-V08B1), Bio-COVID-19_S1 protein_His_Avitag (Acro Biosystems), SARS2-S1-Flag-3Streptag (University of Utrecht), COVID-19_S_protein_RBD_Fc (Acro Biosystems), and SARS2-S1B-2Streptag (University of Utrecht). Proteins were biotinylated by using standard methods.

### Selection of SARS-CoV-2 spike protein-specific DARPin molecules by ribosome display

DARPin libraries(*6*) (N2C and N3C) were used in ribosome display selections(*15,16*) against the SARS-CoV-2 spike protein variants. Four selection rounds were performed per target and per library using decreasing target concentrations and increasing washing stringency (wash buffer composition: 50 mM Tris-HOAc (pH 7.5 at 4°C), 150 mM NaCl, 50 mM Mg(OAc)_2_, 0.05% Tween-20), to increase selection pressure from round 1 to round 4. The number of reverse transcription (RT)-PCR cycles after each selection round was continuously reduced, adjusting to the selection yield due to enrichment of high affinity binders. In detail, the following panning conditions were applied: RD panning round 1: 400 nM target concentration, 8 washes for 1 minutes in wash buffer, 45 PCR cycles; RD panning round 2: 100 nM target concentration, 3 washes for 1 minute in wash buffer, then 3 washes for 15 minutes in wash buffer, followed by 2 washes for 1 minute in wash buffer, 35 PCR cycles; RD panning round 3: 25 nM target concentration, 3 washes for 1 minute in wash buffer, then 3 washes for 30 minutes in wash buffer, followed by 2 washes for 1 minute in wash buffer, 30 PCR cycles; RD panning round 4: 5 nM target concentration, 3 washes for 1 minute in wash buffer, then 3 washes for 45 minutes in wash buffer, followed by 2 washes for 1 minute in wash buffer, 35 PCR cycles (for sub-cloning purposes into expression vector). The 12 resulting pools were then subjected to a binder screening.

### Screening of monovalent DARPin molecules

Monovalent DARPin molecules specifically binding to the S1-RBD, S1-NTD and S2 domains of the spike protein of SARS-CoV-2 in solution were identified by a homogeneous time resolved fluorescence (HTRF) assay using crude extracts of DARPin-expressing *Escherichia coli* (*E. coli*) cells using standard protocols Briefly, DARPin clones selected by ribosome display were cloned into a derivative of the pQE30 (Qiagen) expression vector, *E. coli* XL1-Blue (Stratagene) was transformed and plated on LB-agar (containing 1% glucose and 50 μg/ml ampicillin) and then incubated overnight at 37°C. Single colonies were picked into individual wells of 96 well plates containing 165 μl growth medium (LB containing 1% glucose and 50 μg/ml ampicillin) and incubated overnight at 37°C, shaking at 800 rpm. 150 μl of fresh LB medium containing 50 μg/ml ampicillin was inoculated with 8.5 μl of the overnight culture in a fresh 96-deep-well plate. After incubation for 120 min at 37°C and 850 rpm, expression was induced with IPTG (0.5 mM final concentration) and continued for 6 h. Cells were harvested by centrifugation of the 96-deep-well plates, supernatant was discarded, and the pellets were frozen at -20°C overnight before resuspension in 8.5 μl B-PERII (Thermo Scientific) and incubation for 1 h at room temperature with shaking (600 rpm). Then, 160 μl PBS was added and cell debris was removed by centrifugation (3220 g for 15 min). The extract of each lysed clone was applied as a 1:200 dilution (final concentration) in PBSTB (PBS supplemented with 0.1% Tween 20® and 0.2% (w/v) BSA, pH 7.4) together with 20 nM (final concentration) biotinylated spike protein domain, 1:400 (final concentration) of anti-6His-D2 HTRF antibody – FRET acceptor conjugate (Cisbio) and 1:400 (final concentration) of anti-strep-Tb antibody FRET donor conjugate (Cisbio, France) to a well of a 384-well plate and incubated for 120 min at 4°C. The HTRF was read-out on a Tecan M1000 using a 340 nm excitation wavelength and a 620±10 nm emission filter for background fluorescence detection and a 665±10 nm emission filter to detect the fluorescence signal for specific binding. The extract of each lysed clone was tested for binding to the biotinylated spike protein domains, in order to assess specific binding to the spike protein.

### Cloning of multivalent DARPin molecules

Multivalent DARPin molecules were prepared using Gibson assembly as described(*50*). The individual domains are linked with proline-threonine-rich polypeptide linkers(*50*) flanked by glycine-serine, with a length of 24 amino acids (GSPTPTPTTPTPTPTTPTPTPTGS).

### DAPRin protein production and characterization

DARPin molecules were expressed in *E. coli* and purified as described(*50*). Characterization of monovalent DARPin molecules included SDS-PAGE, size exclusion chromatography, surface plasmon resonance, SARS-CoV-2 pseudotype virus inhibition assay, as well as live virus inhibition assay. Characterization of multivalent DARPin molecules included SDS-PAGE (fully intact size without degradation; not shown), mass spectrometry (expected molecular weight; not shown), size exclusion chromatography coupled to static light scattering, circular dichroism, storage stability (stable at 60°C for 1 week; data not shown), serum stability (stable at 37°C in serum for one week; data not shown), surface plasmon resonance, SARS-CoV-2 pseudotype virus inhibition assay, live virus inhibition assay, hamster pharmacokinetic analysis, and hamster efficacy model as further described below.

### Circular dichroism of DARPin molecules

Circular dichroism measurement was performed with a Jasco J-815 using a 1 cm pathlength cuvette (Hellma) with the monitor sensor inserted in the cuvette. The MRE at 222 nm was followed over a temperature ramp from 20°C to 90°C (heating and cooling). Spectra from 190-250 nm were taken before and after the variable temperature measurement at 20°C. The protein was measured at 0.25 μM in PBS.

### Surface plasmon resonance affinity determination of DARPin molecules

SPR assays were used to determine the binding affinity of monovalent DARPin as well as multivalent DARPin molecules to the spike protein of SARS-CoV-2. All SPR data were generated using a Bio-Rad ProteOn XPR36 instrument with PBS-T (0.005% Tween20) as running buffer. A new neutravidin sensor chip (NLC) was air-initialized and conditioned according to Bio-Rad manual.

Monovalent DARPin molecules: Chemically biotinylated (via lysines) SARS-CoV-2 spike protein 20 (Sino Biologics) was captured to ∼3400 RUs (30 μg/ml, 30 μl/min, 300 s). Two buffer injections (100 μl/min, 60 s) followed by two 12.5 mM NaOH regeneration steps (100 μl/min, 18 s) were applied before the first injections. Mono-domain DARPin proteins were injected (at 50/16.7/5.6/1.9/0.6 nM) for 180 s at 100 μl/min for association and dissociation was recorded for 3600 s (at 100 μl/min). The ligand was regenerated with a 12.5 mM NaOH pulse (100 μl/min, 18 s). The data was double referenced against the empty surface and a buffer injection and fitted according to the 1:1 Langmuir model.

Multivalent DARPin molecules: Chemically biotinylated (via lysines) COVID-19_S_protein_RBD_Fc (Acro Biosystems) was captured to ∼1000 RUs (775 ng/ml, 30 μl/min, 300 s). Two buffer injections (100 μl/min, 60s) followed by two 12.5 mM NaOH regeneration steps (100 μl/min, 18s) were applied before the first injections. One single concentration of 25 nM of each multi-domain drug candidate was injected for 180 s at 100 μl/min for association and dissociation was recorded for 36’000 s (at 100 μl/min). The data was double referenced against the empty surface and a buffer injection. Due to avidity gain, no significant dissociation could be recorded during the measured time.

### Cells and viruses

Vero E6 cells (African green monkey kidney cells, ATCC® CRL1586™) purchased from ATCC (Manassas, VA 20110 USA) were passaged in cell culture medium DMEM (FG0445) containing 10% FBS and supplements (2mM L-Glutamine, Non-essential amino acids and 100 U/ml Penicillin 100 µg/ml Streptomycin and HEPES, all from Biochrom, Berlin, Germany) at 37°C with CO2. SARS-CoV-2 (2019-nCoV/IDF0372/2020) kindly provided by Dr. Sylvie van der Werf from the National Reference Centre for Respiratory Viruses hosted by Institut Pasteur (Paris, France) was propagated in Vero E6 cells in MEM containing 2% FBS and supplements (2%-FBS-MEM) at 37°C and 5% CO_2_.

Virus neutralization capacity of monovalent DARPin and multivalent DARPin molecules was determined for 100 TCID_50_ SARS-CoV-2 by crystal violet staining of protected cells. DARPin molecules were serially diluted from 50 nM to 3.2 pM (in triplicates) in 100 µl cell culture medium (2%-FBS-MEM) enriched with 10 µM human serum albumin (HSA, CSL Behring, Switzerland) (2%-FBS-MEM + HSA) in 96 well plates. The diluted DARPin candidates were exposed to 100 TCID_50_ SARS-CoV-2 (10^4^ TCID_50_/ml) in 100 µl 2%-FBS-MEM + HSA. DARPin/virus mixtures (200 µl) were transferred onto 80% confluent Vero E6 cells. The controls consisted of Vero E6 cells exposed to DARPin molecules only, to determine unspecific effects of the DARPin molecules, of cells exposed to virus suspension only to determine maximal cytopathic effect and of cells incubated with medium only, to determine baseline state of cells. The plates were incubated for 3 days at 37°C and the cytopathic effect determined by staining with 100 µl/well crystal violet solution (spatula tip (∼4 mg) crystal violet powder (Sigma Aldrich) solved in 30 ml 37% formalin and 120 ml PBS (Sigma Aldrich)) for 10 min and washing plates with PBS (Sigma Aldrich). Wells were visually evaluated for complete protection indicated by intact blue/violet cell layer or partial protection in case of >50% intact cell layer.

The effect of neutralization capacity of Multivalent DARPin was evaluated by exposing serial dilutions of the DARPin candidates to increasing titers of SARS-CoV-2 and determining cell protection by CellTiter-Glo assay (Promega, Madison, USA). Serial dilution of DARPin candidates were prepared in 96 well plates in 100 µl cell culture medium (2%-FBS-MEM + HSA) mixed with 100 µl virus suspension of 10^4^ TCID_50_/ml SARS-CoV-2 and incubated for 1 h at 37°C and 5% CO_2_. DARPin/virus mixtures (200 µl) were transferred onto 80% confluent Vero E6 cells and plates incubated at 37°C for 3 days. Cell viability was determined by removing 100 µl supernatant from all wells and adding 100 µl CellTiter-Glo reagent as described in the manufacturers protocol (CellTiter-Glo® Luminescent Cell Viability Assay). Luminescence was read after 2 minutes shaking on an orbital shaker and 10 min incubation at RT using the GloMax instrument (Promega, Madison, USA).

### SARS-CoV-2 VSV pseudotype virus assay

SARS-CoV-2 pseudoviruses were generated as described previously(*51*). A selected panel of 380 DARPin molecules expressed in 96-well format and purified to homogeneity were evaluated for their anti-viral activity in a pseudovirus screening assay. In parallel, 6 monovalent DARPin molecules of known spike domain specificity which were randomly assembled in a set of 192 tri-specific DARPin molecules were included in the screen. For the pseudovirus screening assays, DARPin molecules were diluted in Dulbecco modified Eagle medium (DMEM)-2 % [vol/vol] fetal calf serum (FCS) at the following concentrations: 200 nM, 20 nM, 2nM and 0.2 nM and mixed with an equal volume of DMEM-2 % [vol/vol] FCS containing the VSV-based SARS-CoV-2 pseudoviruses to obtain 2000 IU/well. The mix was incubated for 60 min at 37°C, then inoculated onto Vero E6 cells in a clear bottom white walled 96-well plate during 90 min at 37°C. The inoculum was removed and fresh medium added, and cells further incubated at 37°C for 16 h. Cell were lysed according to the ONE-Glo™ luciferase assay system (Promega, Madison, US) and light emission was recorded using a Berthold® TriStar LB941 luminometer. For the pseudovirus titrations an initial dilution of the drug was followed by two-fold dilutions in quadruplicates in DMEM)-2 % [vol/vol] FCS supplemented with 20 μM human serum albumin (CSL Behring). The mixture was mixed with an equal volume of DMEM-2 % FCS containing SARS-CoV-2 pseudoviruses and incubated for 90 min at 37°C. Neutralizations were performed from 200 to 2000 infectious units (IU) per well, depending on the experiment. Following this incubation, the mix was inoculated onto Vero E6 cells in a clear bottom white walled 96-well plate during 90 min at 37°C. The inoculum was removed and fresh medium added, and cells further incubated at 37°C for 16 h. Cell were lysed according to the ONE-Glo™ luciferase assay system (Promega, Madison, US) and light emission was recorded using a Berthold® TriStar LB941 luminometer. The raw data (relative light unit values) were exported to GraphPad Prism v8.01, and the % neutralization data were normalized to the untreated PsV signal. IC_50_ with 95% confidence interval were estimated by model of nonlinear regression fit with settings for log (inhibitor) vs normalized response curves.

### Laboratory 2: PsV NA for titration curves

Production of VSV pseudotyped with SARS2-S was described previously(*52*). Briefly, HEK-293T cells were transfected with pCAGGS expression vectors encoding MERS-S, SARS-S or SARS2-S carrying a 16-, 28- or 18-a.a. cytoplasmic tail truncation, respectively. One day post transfection, cells were infected with the VSV-G pseudotyped VSVΔG bearing the firefly (Photinus pyralis) luciferase reporter gene. Twenty-four hours later, supernatants containing SARS2-S pseudotyped VSV particles were harvested and titrated on African green monkey kidney Vero E6 (ATCC#CRL-1586) cells.

In the virus neutralization assay, DARPin molecules were threefold serially diluted at two times the desired final concentration in DMEM supplemented with 10 μM human serum albumin (CSL Behring), 100 U/ml Penicillin and 100 µg/ml Streptomycin (Lonza). Monoclonal antibodies against MERS-S (2), SARS-S or SARS2-S were included as a positive control(*52*). Diluted DARPin molecules and mAbs were incubated with an equal volume of pseudotyped VSV particles for 1 hour at room temperature, inoculated on confluent Vero E6 monolayers in 96-well plate, and further incubated at 37 °C for 24 hours. Cells were lysed with Luciferase Cell Culture Lysis 5X Reagent (Promega) at room temperature for 30 min. Luciferase activity was measured on a Berthold Centro LB 960 plate luminometer using D-luciferin as a substrate (Promega). The half maximal inhibitory concentrations (IC_50_) were determined using 4-parameter logistic regression (GraphPad Prism version 8).

### Cryo-electron microscopy

4 μl of purified S ectodomain (9 μM) was mixed with 1 μl of 50 μM mono-DAPRin #3, #9 or #10, and incubated for 15 seconds at room temperature. 3 μl of sample was then dispensed on Quantifoil R1.2/1.3 200-mesh grids (Quantifoil Micro Tools GmbH) that had been freshly glow discharged for 30 s at 20 mA. Grids were blotted using blot force +2, for 5 s using Whatman No. 1 filter paper and immediately plunge-frozen into liquid ethane cooled by liquid nitrogen using a Vitrobot Mark IV plunger (Thermo Fisher Scientific) equilibrated to ∼95% relative humidity, 4°C. Movies of frozen-hydrated specimens were collected using Glacios Cryo-TEM (Thermo Fisher Scientific) operating at 200 keV and equipped with a Falcon 4 Direct Electron Detector (Thermo Fisher Scientific). For additional analysis of monovalent DARPin #3, 4 μl of purified S ectodomain (18 μM) was mixed with 1 μl of 100 μM DARPin, and incubated for 60 s at room temperature. Grids were prepared as described above, and movies were collected using a Titan Krios Cryo-TEM (Thermo Fisher Scientific) operating at 300 keV and equipped with a Falcon 4 Direct Electron Detector (Thermo Fisher Scientific). All cryo-EM data were acquired using the EPU 2 software (Thermo Fisher Scientific) with a 30-degree stage tilt to account for preferred orientation of the samples. Movies were collected in electron counting mode at 92,000x (Glacios) or 75,000x (Titan Krios), corresponding to a pixel size of 1.1 Å/pix or 1.045 Å/pix over a defocus range of -1.25 to -2.5 μm.

### Image processing

Movie stacks were manually inspected and then imported in Relion version 3.1(*53*). Drift and gain correction were performed with MotionCor2(*54*), and GCTF(*55*) was used to estimate the contrast transfer function for each movie. Particles were automatically picked using the Laplacian-of-Gaussian (LoG) algorithm and then Fourier binned (2 x 2) particles were extracted in a 160-pixel box. The extracted particles were subjected to two rounds of 2D classification, ignoring CTFs until the first peak. Using the ‘molmap’ command in UCSF chimera(*56*), a SARS-CoV-2 spike structure (PDB ID: 6VSB)(*23*) was used to generate a 50 Å resolution starting model for 3D classification. Particles selected from 2D classification were subject to a single round of 3D classification (with C1 symmetry). Particles belonging to the best classes were re-extracted unbinned in a 320-pixel box, 3D auto-refined (with C1 or C3 symmetry) and post-processed. Iterative rounds of per particle defocus estimation, 3D auto-refinement and post-processing were used to account for the 30-degree stage tilt used during data collection. When CTF refinement did not yield any further improvement in resolution, Relion’s Bayesian polishing procedure was performed on the particle stacks, with all movie frames included, followed by 3D auto-refinement and post-processing. Subsequently, additional rounds of per particle defocus estimation, 3D auto-refinement and post-processing were performed on the polished particles until no further improvement in resolution or map quality was observed. The nominal resolution for each map was determined according to the ‘gold standard’ Fourier shell correlation (FSC) criterion (FSC = 0.143) and local resolution estimations were performed using Relion. Map sharpening was performed using DeepEMhancer(*57*) as implemented in COSMIC2(*58*). To improve the quality of the mono-DARPin #3 density in the fully open spike reconstruction, a focused 3D classification approach was employed. Briefly, each particle contributing to the final C3-symmetry–imposed reconstruction was assigned three orientations corresponding to its symmetry related views using the “relion_particle_symmetry_expand” tool. A soft mask was placed over the map to isolate the mono-DARPin #3-bound RBD, and the symmetry-expanded particles were subjected to masked 3D classification without alignment using a regularization parameter (‘T’ number) of 20. Particles corresponding to the 3D class with the best resolved DARPin density were re-extracted in a 200-pixel box and centered on the mask used for focused classification. In conjunction with this, the signal for the protein outside the masked was subtracted. The re-extracted particles were then 3D auto-refined (with C1 symmetry) using local angular searches (1.8 degrees) and sharpened using DeepEMhancer(*57*). Three copies of the locally refined map were aligned to the globally refined map using the UCSF Chimera ‘fit in map’ tool and resampled using the ‘vop resample’ command. Finally, a composite map was generated using the “vop add” command. An overview of the image processing workflows for each of the monovalent DARPin samples is shown in Supplementary Figure 2.

### Molecular modeling of mono and multivalent DARPin molecules

Homology models of monovalent DARPin molecules #3 and #9 were generated with Rosetta(*59*). The consensus designed ankyrin repeat domain PDB ID:2xee was used as template. Mutations were introduced with RosettaRemodel(*60*) with fixed backbone, and the structure was refined with RosettaRelax(*61*). Forty refined structures were clustered using RosettaCluster with 0.3 Å radius, and the lowest-energy model from the largest cluster served as the final model. These models were then used for fitting domain #3 and #9 into the observed electron density generated from the complex structure of the spike protein.

To facilitate accurate fitting of the DARPin coordinates into their respective cryo-EM maps, difference density maps of the bound DARPin molecules were produced as described previously(*62*). For monovalent DARPin #3, the atomic coordinates of a fully-open spike ectodomain (PDB ID: 6XCN) were fitted into the EM density using the UCSF Chimera ‘fit in map’ tool. The Fab component of the model was deleted, and then the ‘molmap’ command was used to simulate a 7 Å resolution density map. This simulated map was then resampled on the grid of the experimental cryo-EM density map using the ‘vop resample’ command. The ‘vop subtract’ command was then used to subtract the value of the simulated map from the experimental map. The ‘minRMS’ option was used to automatically scale the simulated map to minimize the root-mean-square sum of the resulting subtracted values at grid points within the lowest contour of simulated map. The UCSF Chimera ‘fit in map’ tool was then used to fit monovalent DARPin #3 into the difference density until the correlation between the map and model did not improve any further, and ensuring that the epitope binding surface of the monovalent DARPin was orientated towards the spike ectodomain. This workflow was then repeated for monovalent DARPin #9, using the fully closed spike coordinates (PDB ID: 6ZGE).

The PDB file with the coordinates of the trimer of domain #3:RBD was used as an input structure for the conceptual modeling of MR-DC bound to the spike ectodomain as shown in Supplementary Figure 8. In both models, the open RBD domain from PDB ID 6vyb was used to generate three RBDs in the open conformation(*13*). For the conceptual modeling of MM-DC a similar approach was used by fitting domain #9 into the observed density of the NTD cryo-EM structure. Additionally, a structurally resolved NTD domain from PDB 7c2l was used(*26*), and a binding domain model of S2 (#10) was placed manually to a potential interacting site on S2 (non-glycosylated region within reasonable distance from the binding domain #9, based on linker length). For both structures, HSA binding DARPin models were placed and the linkers between each binding domain were modeled using Rosetta modeling tools(*60*).

In a last step, the models of monovalent DARPin #3:RBD (residues 303-526) and monovalent DARPin #9:NTD (residues 14-303) were refined with Rosetta. The structures were pre-relaxed for docking and served as input for local, high-resolution docking with RosettaDock(*63*) with fixed backbone. Five hundred models were generated and clustered with 1 Å radius (RosettaCluster). Two largest clusters were inspected and the lowest-energy model from more conserved group (i.e., with lower rigid-body perturbation from the input structure) was taken further for additional all-atom refinement with RosettaRelax(*61*), with protocol optimized for interfaces (InterfaceRelax2019). Fifty models were generated, and the lowest scoring model was selected. This model was used to describe the interactions between DARPin molecules and their target domains. Figures were generated using LigPlot(*64*), UCSF Chimera(*56*), UCSF ChimeraX(*65,66*), PyMOL (The PyMOL Molecular Graphics System, Version 2.0, Schrödinger, LLC) and BioRender (BioRender.com).

Prophylactic Syrian golden hamster model for the assessment of antiviral potency of candidate MR-DC The study was performed at Viroclinics Xplore, Schaijk, The Netherlands.

### Virus used for Syrian golden hamster study

SARS-CoV-2 isolate BetaCoV/Munich/BavPat1/2020 was kindly provided by Prof. Dr. C. Drosten (European Virus Archive Global # 026V-03883). With a history of 1 passage in Vero-TMPRSS2 and 3 passages in Vero E6 cells (ATCC), the seed stock was titrated in Vero E6 cells to a concentration of 7.1 log10 TCID_50_/ml. The seed stock was thawed and diluted in cold phosphate-buffered saline (PBS) prior to infection.

### Experimental design

Twenty-four specific-pathogen-free (SPF) 15 weeks-old Syrian golden hamsters (Mesocricetus auratus, females and males, approximate body weights of 160g, provided by Envigo) were uniquely identified using individually-coded animal markers. They were housed in elongated type 2 group cages with two animals per cage under BSL-III conditions during the experiment. They were kept according to the standards of Dutch law for animal experimentation and were checked daily for overt signs of disease. The study was carried out following approval by an independent animal welfare body (approval AVD277002015283-WP13) and complied with all relevant ethical regulations for animal testing.

Four groups of six hamsters were treated with multivalent DARPin molecule MR-DC via the intraperitoneal route with 16 µg, 160 µg, or 1600 µg doses per animal or with a placebo 24 hours prior to infection and subsequently animals were inoculated intra-nasally with 100 µL PBS containing 5×10^4^ TCID_50_ SARS-CoV-2. The inoculum was instilled dropwise using a pipette and equally divided over both nostrils. The animals were weighed regularly and throat swabs, for quantitative PCR and infectious virus titration, were collected on a daily basis. For all animal procedures, the animals were sedated with isoflurane (3-4%/O2).

Upon necropsy at day 4 post infection, full-body gross pathology was performed for each animal and all abnormalities recorded and described. All lung lobes were inspected, and the percentage of affected lung tissue was estimated by eye. Samples of the left nasal turbinates, trachea and the entire left lung (often with presence of the primary bronchi) were preserved in 10% neutral buffered formalin for histopathology. Samples of the right lung parenchyma and right nasal turbinates were frozen for quantitative PCR and virus titration.

### Virology

Throat swabs and homogenized tissue samples (lungs and nasal turbinates) were thawed and tested for the presence of SARS-CoV-2 infectious virus using virus titration. To this end, quadruplicate 10-fold serial dilutions were used to determine the virus titers on confluent layers of Vero E6 cells. Serial dilutions of the samples (throat swabs and tissue homogenates) were prepared and incubated on Vero E6 monolayers for 1 hour at 37°C. Vero E6 monolayers were washed and incubated with infection medium for 4-6 days at 37°C after which plates were scored for cytopathogenic effect (CPE) using the vitality marker WST-8. Viral titers (TCID_50_) were calculated using the method of Spearman-Karber.

### Histopathology

Samples of tissue from the respiratory tract (lung, trachea, and nasal turbinates) and lymphoid organs (spleen and mandibular lymph node) were fixed in 10% neutral buffered formalin (24-48h) and embedded in paraffin. Tissue sections were then stained with hematoxylin and eosin and examined by light microscopy. A semi-quantitative histopathological analysis was performed, and findings scored using the following grades: absent (grade 0), minimal (grade 1), mild (grade 2), moderate (grade 3) or marked (grade 4). The average score for the dose groups was calculated for each finding in the respiratory tract (Figure 4b) and lymphoid organs (Figure 4c).

### Hamster pharmacokinetic study

Single-dose intraperitoneally administered dose pharmacokinetic measurements in female hamsters (n = 6 per group) were performed at 1 mg/kg and 10 mg/kg. Blood samples were collected pre-dose and again at 1 h, 4 h, 8h, 12h, 24 h, 48 h, 72 h, 96 h and 168 h post-injection. Serum concentrations were determined by sandwich ELISA using RBD as capture reagent and an anti-His-tag antibody as detection reagent and using a standard curve. Pharmacokinetic parameters were determined using the software Phoenix WinNonLin (Certara, Princeton, USA) or GraphPadPrism (GraphPad Software, La Jolla,USA) and non-compartmental analyses.

**Supplementary Figure 1:**
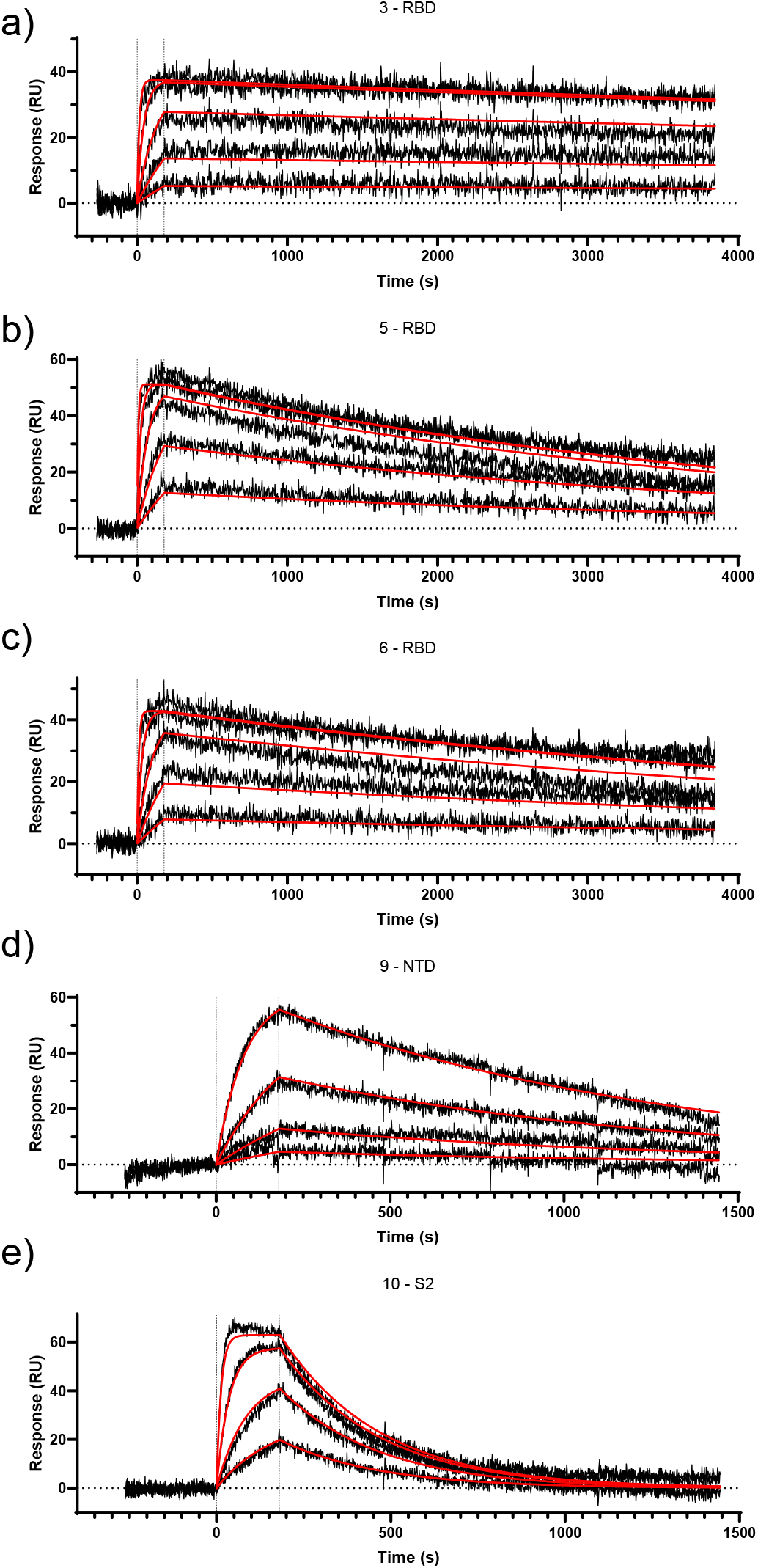
Surface plasmon resonance (SPR) sensorgrams of the monovalent DARPin molecules #3 (a), #5 (b), #6 (c), #9 (d), #10 (e), incorporated in MR-DC (a-c) and MM-DC (c-e) binding to immobilized trimeric spike protein. DARPin concentrations for a-c: 50/16.67/5.56/1.85/0.62 nM. DARPin concentrations for (d) and (e): 16.67/5.56/1.85/0.62 nM. Affinity values of monovalent DARPin molecules are listed in **Table 1**.

**Supplementary Figure 2:**
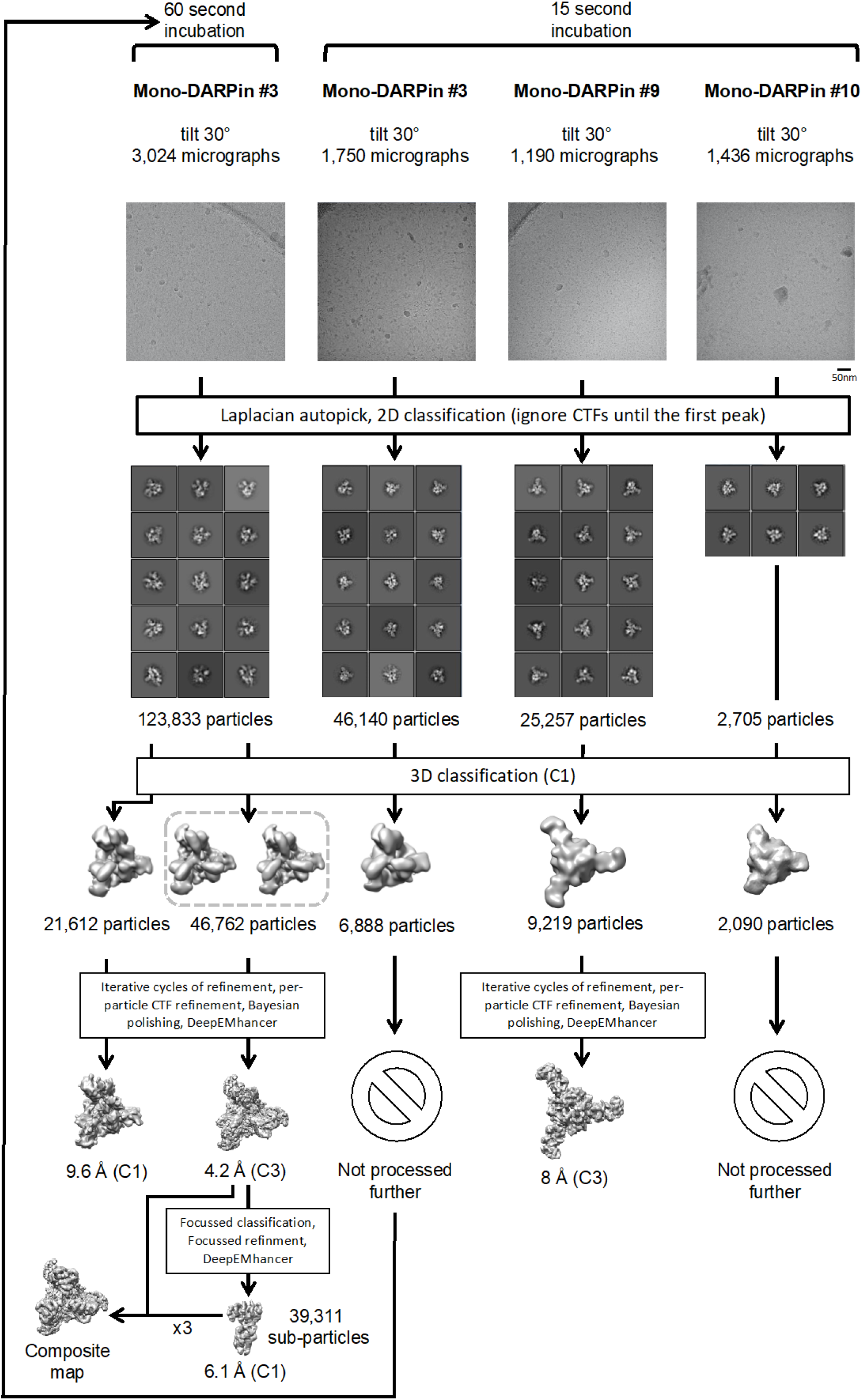
Single-particle cryo-EM image processing workflows for each of the monovalent DARPin samples.

**Supplementary Figure 3:**
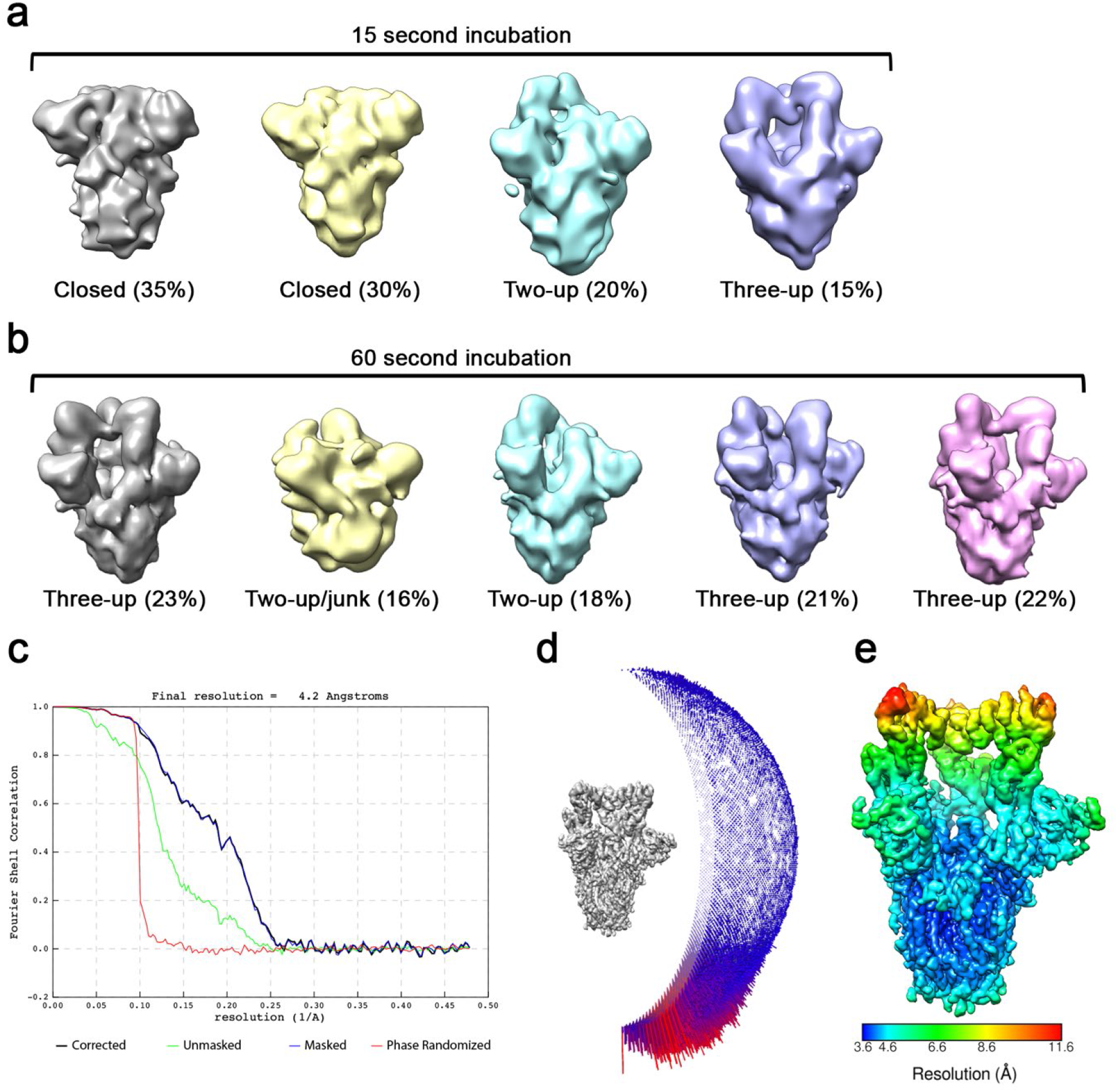
a) 3D classes obtained from spike ectodomains incubated with monovalent DARPin #3 for 15 seconds, and b) for 60 seconds. c) Gold-standard Fourier shell correlation (FSC) curve generated from the independent half maps contributing to the 4.2 Å resolution density map. d) Angular distribution plot of the final C3 refined EM density map. e) The EM density map of the spike ectodomain bound to three copies of monovalent DARPin #3, colored according to local resolution.

**Supplementary Figure 4:**
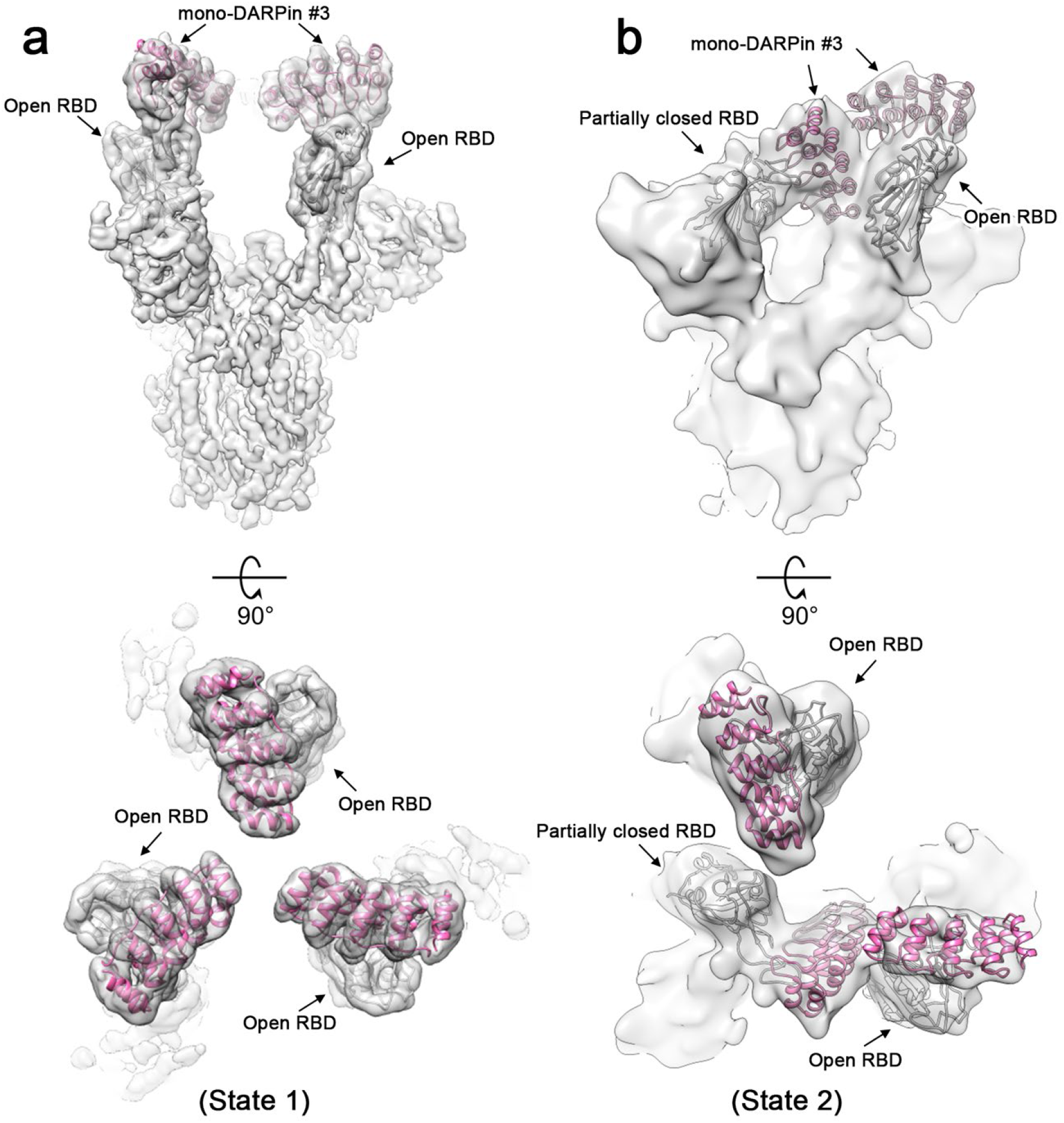
a) Cryo-EM density for state 1 and b) state 2 of the SARS-CoV-2 spike ectodomain in complex with the RBD targeting monovalent DARPin #3, shown as two orthogonal views. The pseudo-atomic model of monovalent DARPin #3 in complex with RBD, derived from molecular docking experiments, is fitted in each of the spike protomers and colored grey and pink, respectively.

**Supplementary Figure 5:**
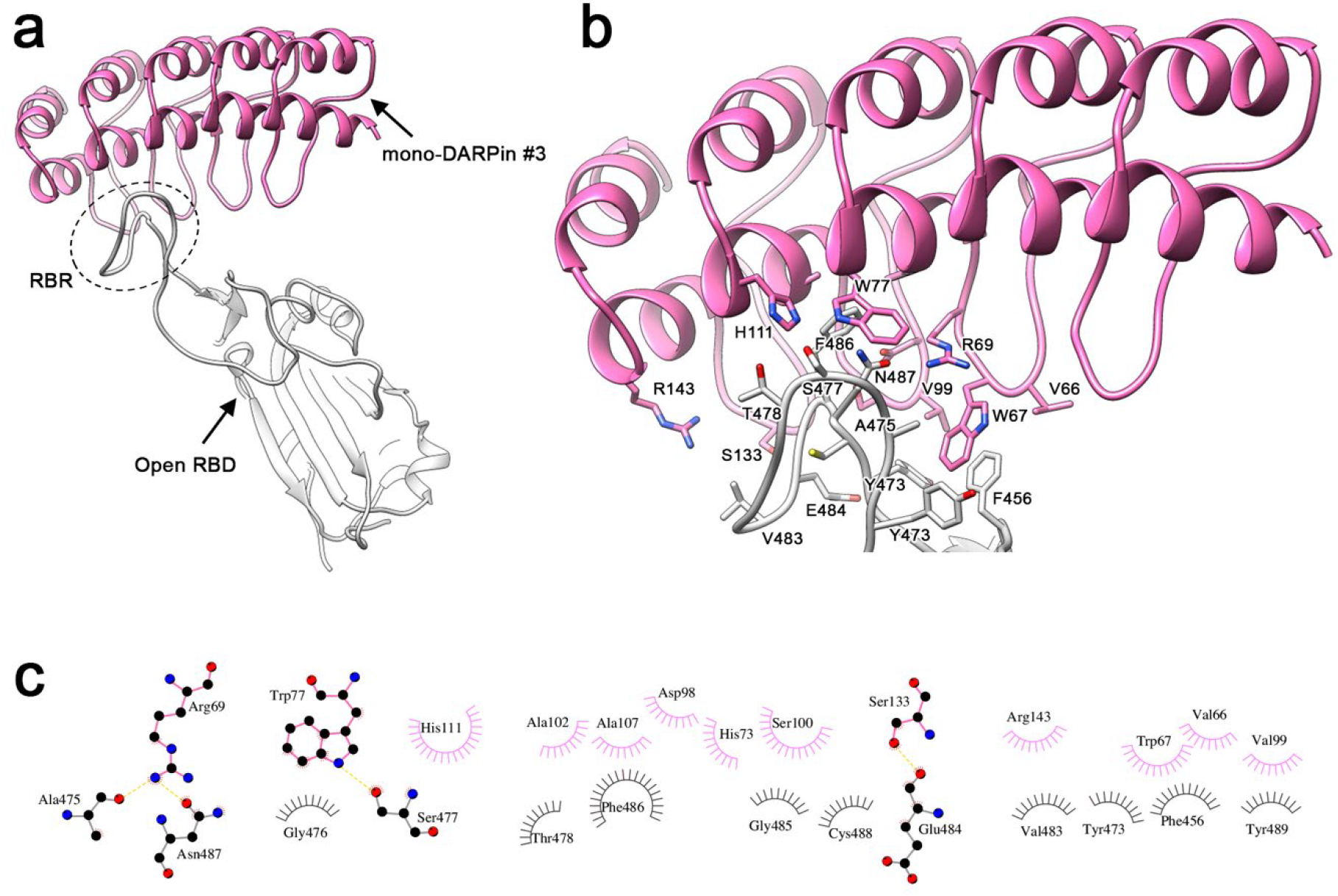
a) Pseudo-atomic model of monovalent DARPin #3 in complex with the SARS-COV-2 spike RBD colored grey and pink, respectively. b) Zoomed in view of the interface between monovalent DARPin #3 and RBD. c) DimPlot(*67*) representation of putative interacting residues between the spike ectodomain RBD and monovalent DARPin #3, identified through molecular docking experiments. Residues participating in hydrophobic interactions are shown as spoke arcs. Residues participating in hydrogen bonding are shown as sticks, and hydrogen bonds are shown as yellow dotted lines. Residues are coloured grey and pink for spike and monoDARPin #3, respectively.

**Supplementary Figure 6.**
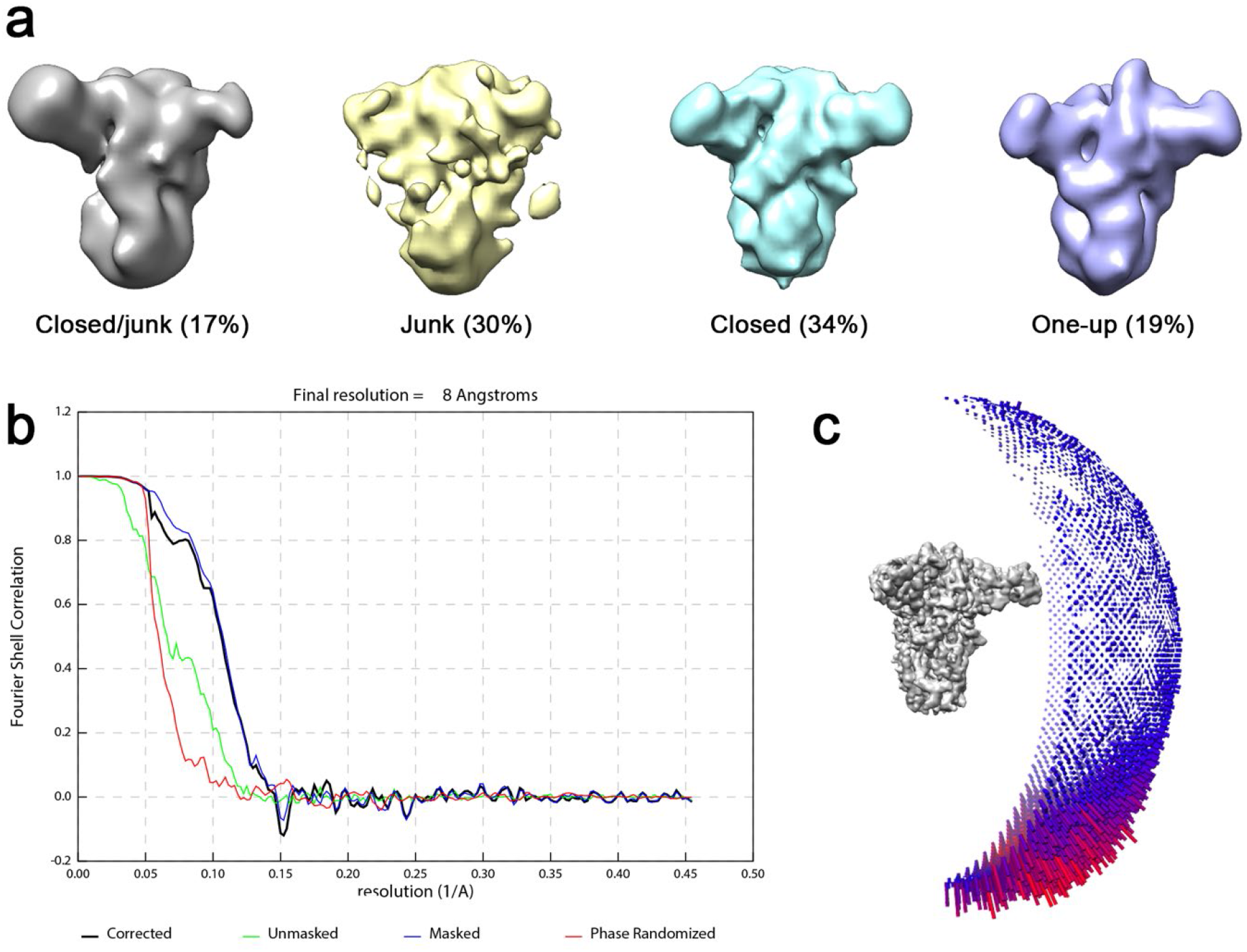
a) 3D classes obtained from spike ectodomains incubated with monovalent DARPin #9 for 15 seconds. b) Gold-standard Fourier shell correlation (FSC) curve generated from the independent half maps contributing to the 8 Å resolution density map. c) Angular distribution plot of the final C3 refined EM density map.

**Supplementary Figure 7:**
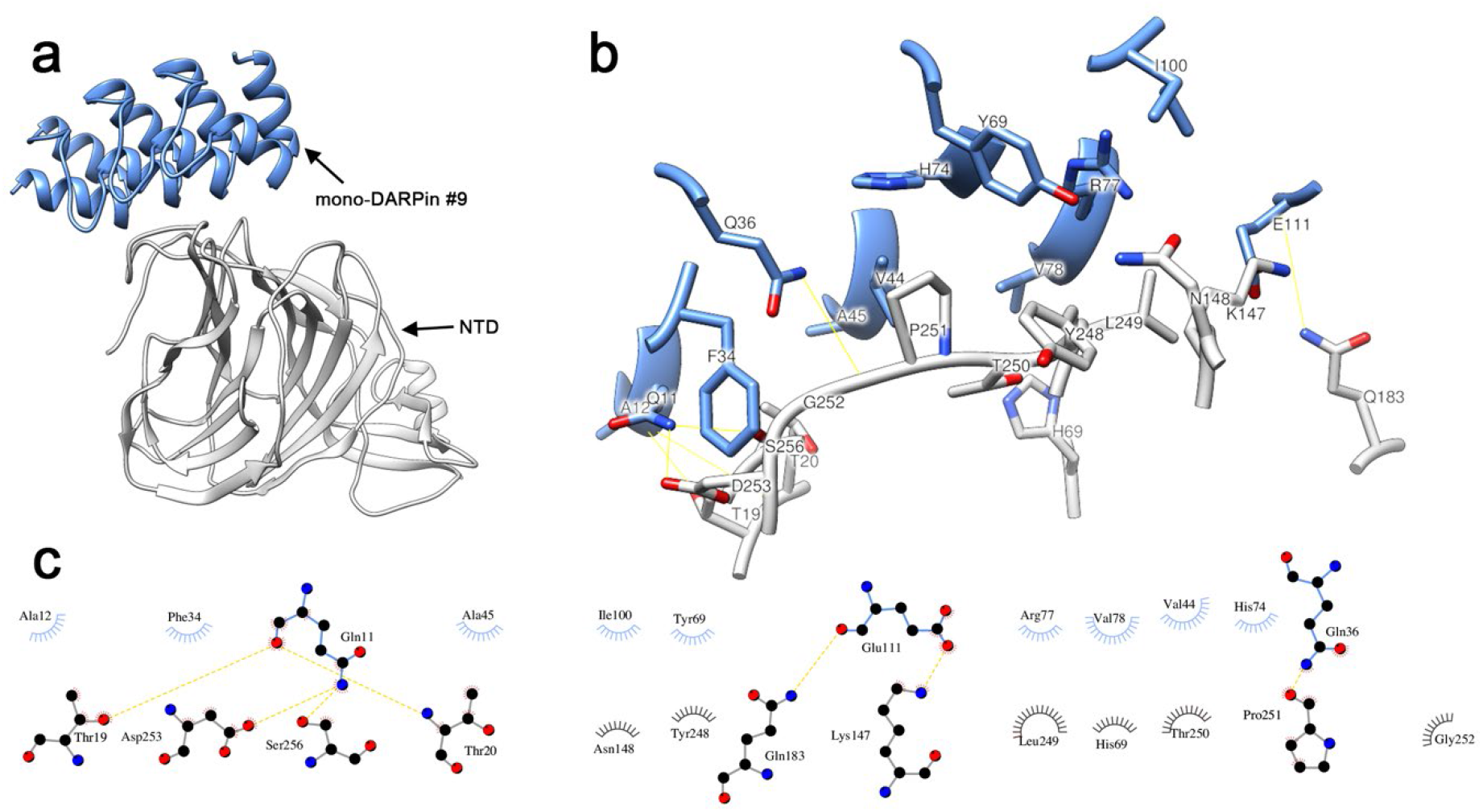
a) Pseudo-atomic model of monovalent DARPin #9 in complex with the SARS-COV-2 spike NTD colored blue and grey, respectively. b) Zoomed in view of the interface between monovalent DARPin #9 and NTD. c) DimPlot(*67*) representation of putative interacting residues between the spike ectodomain NTD and monovalent DARPin #9, identified through molecular docking experiments. Residues participating in hydrophobic interactions are shown as spoke arcs. Residues participating in hydrogen bonding are shown as sticks, and hydrogen bonds are shown as yellow dotted lines. Residues are coloured grey and blue for spike and monovalent DARPin #9, respectively.

**Supplementary Figure 8:**
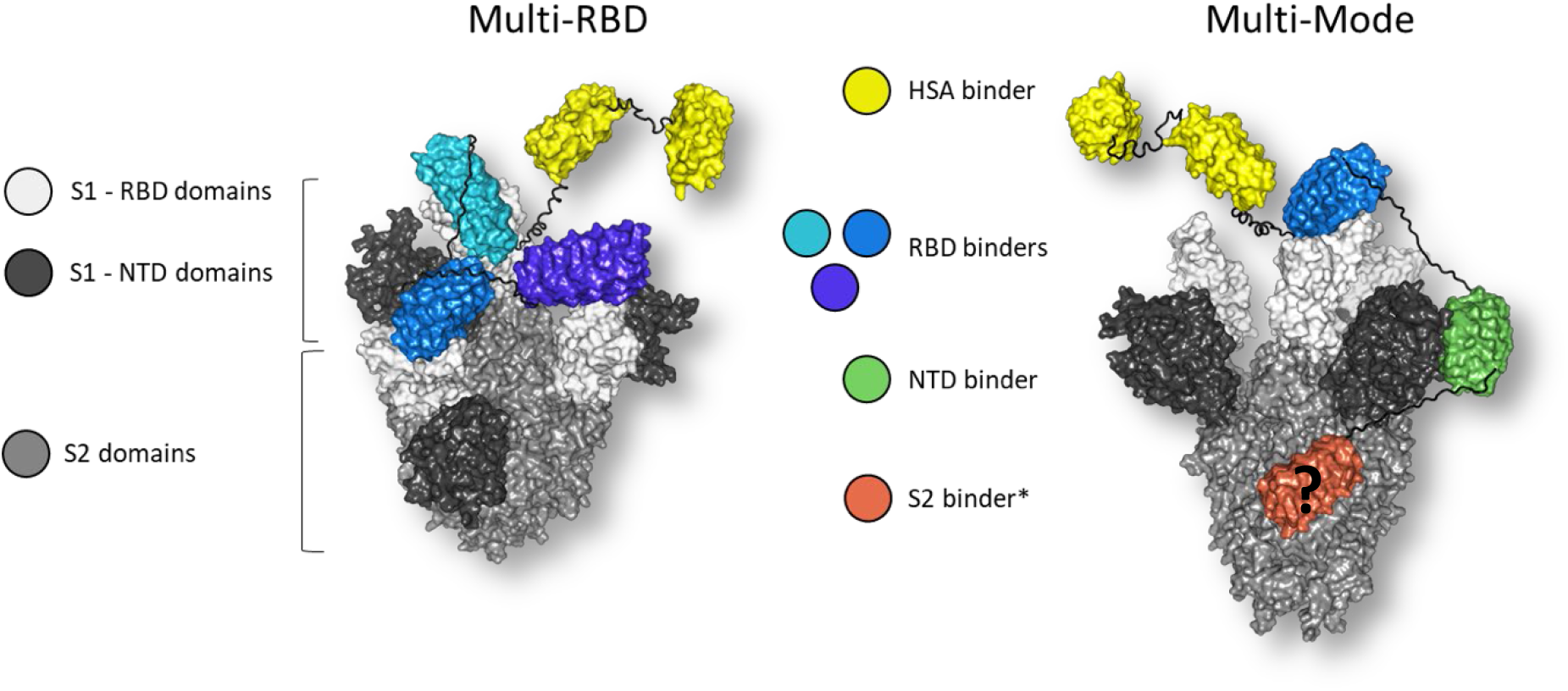
Molecular model of multivalent DARPin molecule MM-DC (right), consisting of five DARPin domains (yellow: HSA-binding domains, blue: RBD-binding domain, green: NTD-binding domain, orange: S2-binding domain) bound to the spike ectodomain. Linkers are shown in black. Molecular model of MR-DC (left) consisting of five DARPin domains (yellow: HSA-binding domains, shades of blue: RBD-binding domains) bound to the RBDs (white) of the spike ectodomain (grey). Linkers are shown in black. Position of RBD and NTD binders guided by Cryo-EM data (Figure 3) - *For the S2 binder, the epitope is unknown.

**Supplementary Figure 9:**
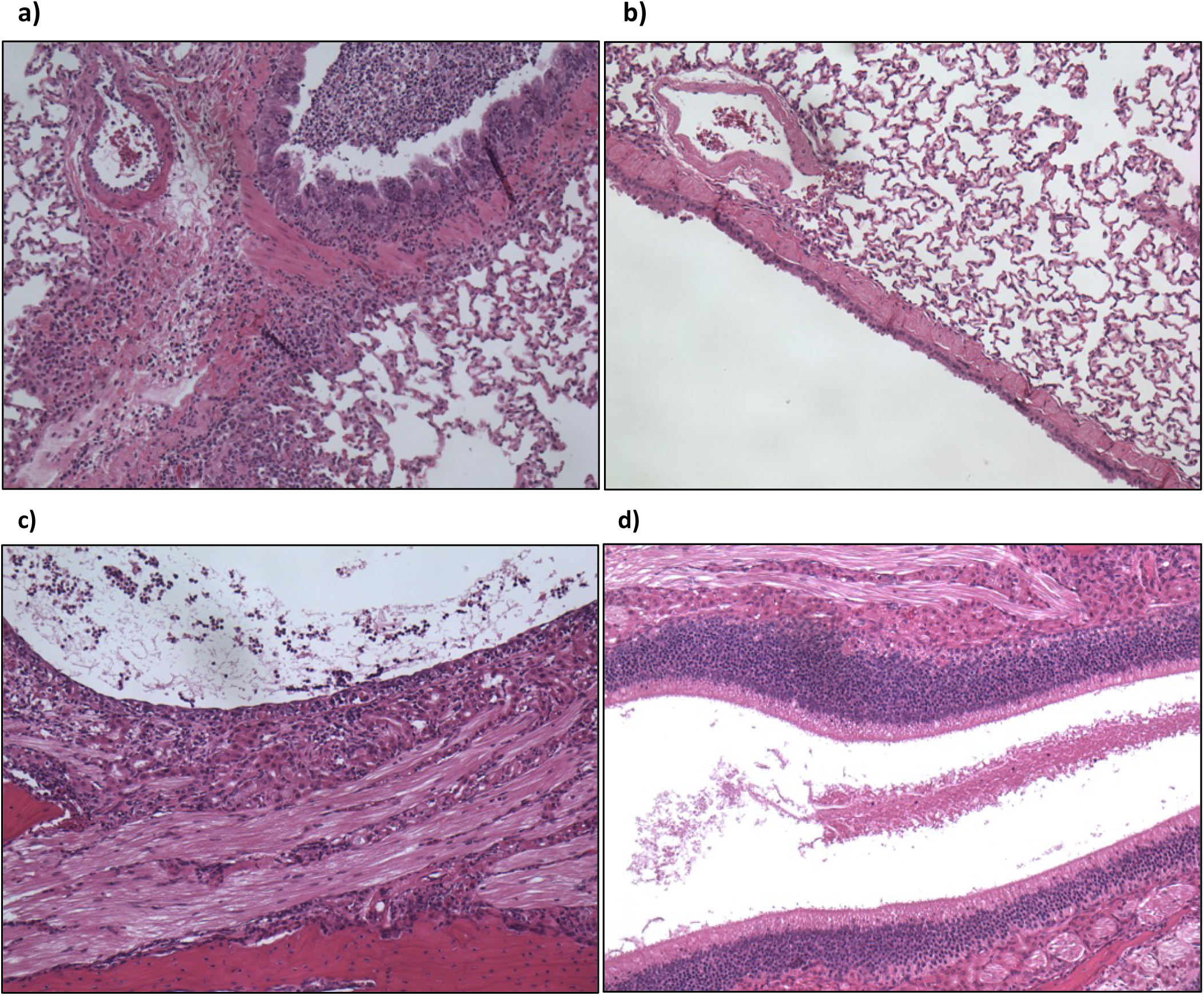
Representative histopathological microscopic images of lung and nasal turbinates on day 4. a) Lung from a hamster treated with vehicle: Moderate mixed inflammatory cell infiltrate (lymphocytes, macrophages, plasma, cells and granulocytes) of the bronchial/bronchiolar epithelium and underlying lamina propria extending to the adjacent blood vessels, alveolar interstitium and spaces with prominent Type II alveolar pneumocytes. Some degeneration, regeneration and disorganization of the bronchial epithelium with single cell necrosis is also evident. b) Lung from a hamster treated with 1600 ug: Very minimal mixed inflammatory cell infiltrate (lymphocytes, macrophages, plasma, cells and granulocytes) of the bronchial/bronchiolar epithelium and underlying lamina propria. c) Nasal turbinates from a hamster treated with vehicle: A marked mixed inflammatory cell infiltrate (lymphocytes, macrophages, plasma, cells and granulocytes) of the olfactory epithelium with degeneration, regeneration and disorganization of the epithelium, single cell necrosis and inflammatory exudate in the lumen of the nasal cavity. d) Nasal turbinates from a hamster treated with 1600 ug: A mild mixed inflammatory cell infiltrate (lymphocytes, macrophages, plasma, cells and granulocytes) of the olfactory epithelium with degeneration, regeneration and disorganization of the epithelium, single cell necrosis and inflammatory exudate in the nasal cavity.

**Supplementary Figure 10:**
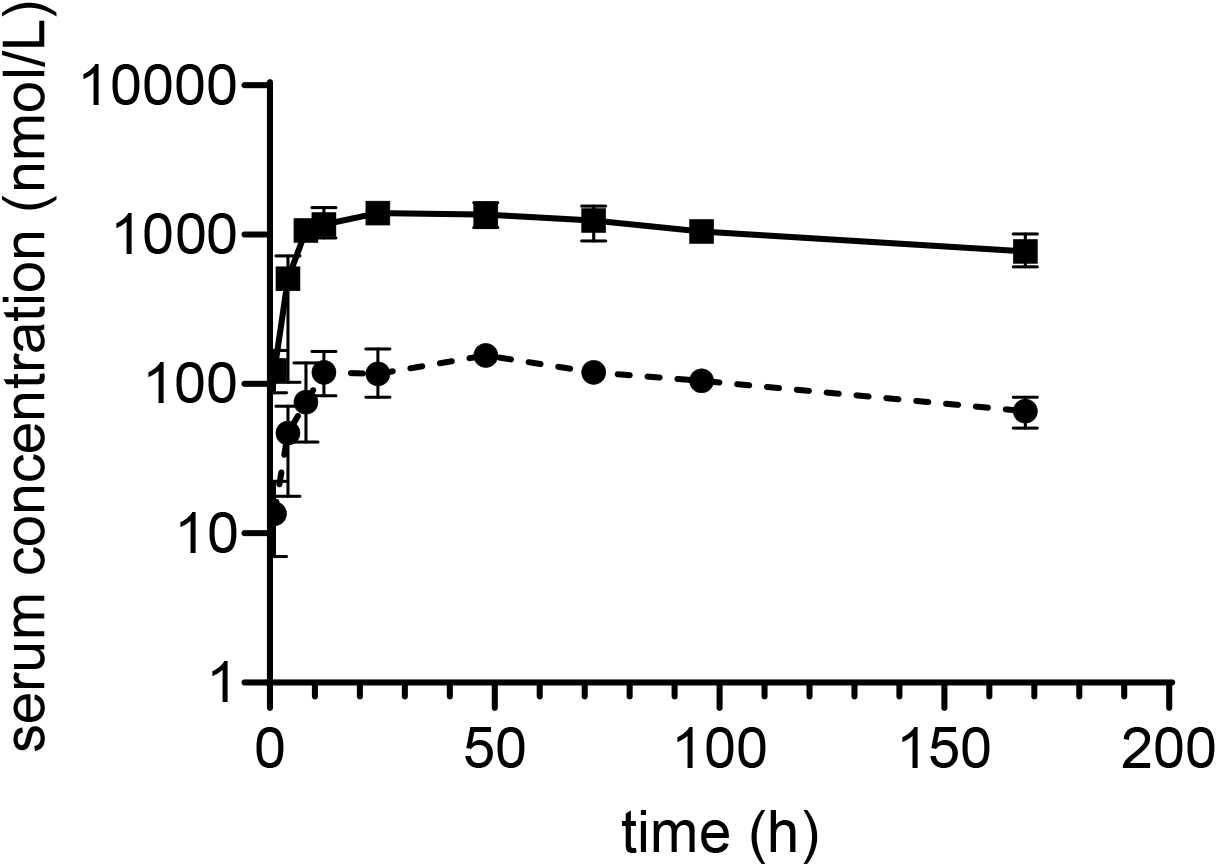
Pharmacokinetic analysis of MR-DC exposure at concentrations of 1 mg/kg (dashed line) and 10 mg/kg (solid line), following i.p. administration. Elimination half-life was calculated to be 4-6 days.

**Supplementary Table 1:**
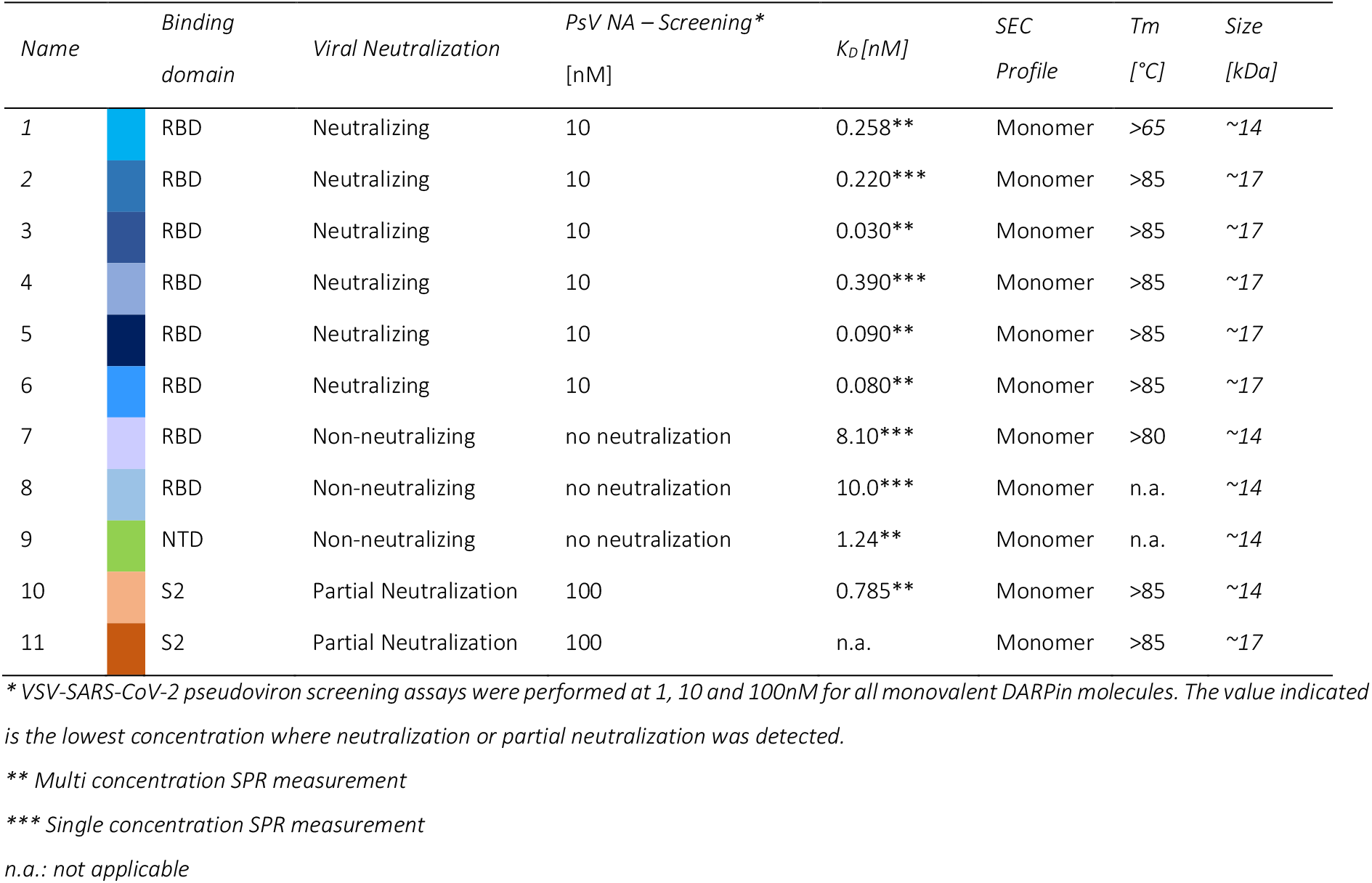
List of spike protein-binding monovalent DARPin molecules and their properties

| Name | Binding domain | Viral Neutralization | PsV NA – Screening*<br>[nM] | K<sub>D</sub> [nM] | SEC Profile | T<sub>m</sub> [°C] | Size [kDa] |
| --- | --- | --- | --- | --- | --- | --- | --- |
| 1 | RBD | Neutralizing | 10 | 0.258** | Monomer | >65 | ~14 |
| 2 | RBD | Neutralizing | 10 | 0.220*** | Monomer | >85 | ~17 |
| 3 | RBD | Neutralizing | 10 | 0.030** | Monomer | >85 | ~17 |
| 4 | RBD | Neutralizing | 10 | 0.390*** | Monomer | >85 | ~17 |
| 5 | RBD | Neutralizing | 10 | 0.090** | Monomer | >85 | ~17 |
| 6 | RBD | Neutralizing | 10 | 0.080** | Monomer | >85 | ~17 |
| 7 | RBD | Non-neutralizing | no neutralization | 8.10*** | Monomer | >80 | ~14 |
| 8 | RBD | Non-neutralizing | no neutralization | 10.0*** | Monomer | n.a. | ~14 |
| 9 | NTD | Non-neutralizing | no neutralization | 1.24** | Monomer | n.a. | ~14 |
| 10 | S2 | Partial Neutralization | 100 | 0.785** | Monomer | >85 | ~14 |
| 11 | S2 | Partial Neutralization | 100 | n.a. | Monomer | >85 | ~17 |

\* VSV-SARS-CoV-2 pseudovirus screening assays were performed at 1, 10 and 100nM for all monovalent DARPins. The value indicated is the lowest concentration where neutralization or partial neutralization was detected.
\*\* Multi concentration SPR measurement
\*\*\* Single concentration SPR measurement
n.a.: not applicable

**Supplementary Table 2:**
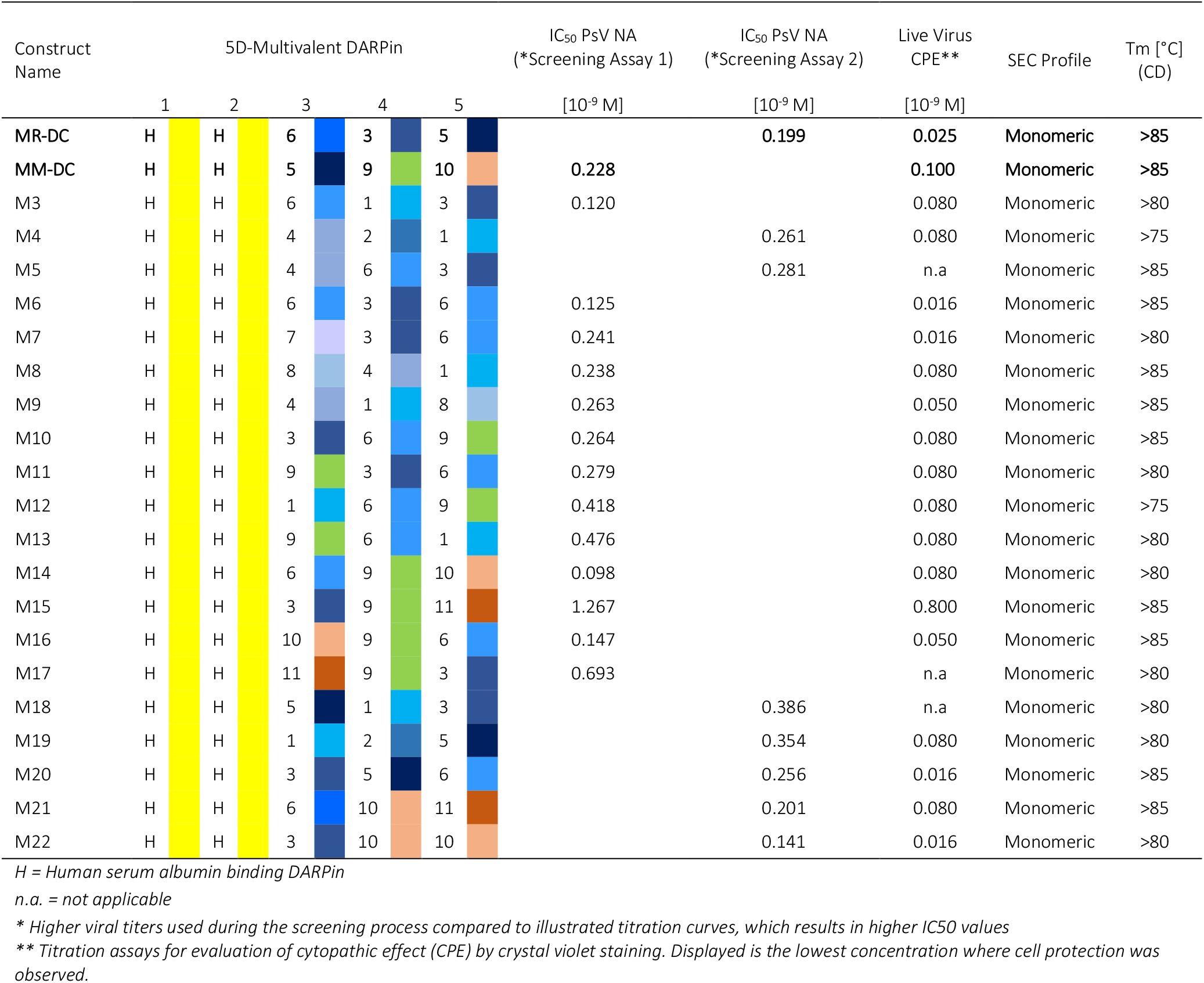
Properties of SARS-CoV-2 inhibiting DARPin candidates

**Supplementary Table 3:** Neutralization potency (IC_50_, [pM]) for multivalent DARPin molecules MR-DC and MM-DC for SARS-CoV-2 spike protein variants, frequently observed in sequencing data of globally appearing serotypes, were evaluated by PsV NA.

|  | wt | G476S | V483A | D614G | D614G x Q675H |
| --- | --- | --- | --- | --- | --- |
| MR-DC | 16.53 | 27.08 | 27.48 | 11.77 | 12.11 |
| MM-DC | 5.48 | 14.46 | 32.40 | 4.64 | 22.44 |

**Supplementary Table 4:** Cryo-EM data collection and image processing information.

| Monovalent DARPIn (no.) | #3 | #3 | #3 | #9 | #10 |
| --- | --- | --- | --- | --- | --- |
| Magnification | 75,000 | 75,000 | 92,000 | 92,000 | 92,000 |
| Voltage (kV) | 300 | 300 | 200 | 200 | 200 |
| Electron exposure (e-/Å²) | 40 | 40 | 40 | 40 | 40 |
| Defocus range (µm) | 1.25-2.5 | 1.25-2.5 | 1.25-2.5 | 1.25-2.5 | 1.25-2.5 |
| Pixel size (Å) | 1.045 | 1.045 | 1.1 | 1.1 | 1.1 |
| Symmetry imposed | C3 | C1 | N/A | C3 | N/A |
| Initial particle images (no.) | 123,833 | 123,833 | 46,140 | 25,257 | 2,705 |
| Final particle images (no.) | 46,762 | 21,612 | 6,888 | 9,219 | 2,090 |
| Map resolution (Å) | 4.2 | 9.6 | N/A | 8 | N/A |
| FSC threshold | 0.143 | 0.143 | N/A | 0.143 | N/A |
| Map resolution range (Å) | 3.7-14.1 | 8.2-26 | N/A | 6.9-18.5 | N/A |

**Supplementary Table 5:** Overview of fermentation runs performed with anti-SARS-Cov-2 multivalent DARPin molecule MP0420 at different scales. Expression yields presented in gram product per liter fermentation broth were determined by SDS-PAGE.

| Run | Status | Fermenter Scale | Harvested Amount | Expression Yield |
| --- | --- | --- | --- | --- |
| 1 | Development | 5 L | 4.9 kg | 12.6 g/L |
| 2 | Development | 5 L | 4.9 kg | 12.5 g/L |
| 3 | Development | 5 L | 4.9 kg | 11.1 g/L |
| 4 | GMP | 100 L | 101 kg | 11.3 g/L |
| 5 | GMP | 100 L | 101 kg | 12.3 g/L |

## Notes

### Summary of Updates

This version of the manuscript has been revised to update Figures and add additional data (e.g. Fig 3).

